# Multi-color fluorescence live-cell imaging in *Dictyostelium discoideum*

**DOI:** 10.1101/2024.10.22.619593

**Authors:** Hidenori Hashimura, Satoshi Kuwana, Hibiki Nakagwa, Kenichi Abe, Tomoko Adachi, Toyoko Sugita, Shoko Fujishiro, Gen Honda, Satoshi Sawai

## Abstract

The cellular slime mold *Dictyostelium discoideum*, a member of the Amoebozoa, has been extensively studied in cell and developmental biology. *D. discoideum* is unique in that they are genetically tractable, with a wealth of data accumulated over half a century of research. Fluorescence live-cell imaging of *D. discoideum* has greatly facilitated studies on fundamental topics, including cytokinesis, phagocytosis, and cell migration. Additionally, its unique life cycle places *Dictyostelium* at the forefront of understanding aggregative multicellularity, a recurring evolutionary trait found across the Opisthokonta and Amoebozoa clades. The use of multiple fluorescent proteins (FP) and labels with separable spectral properties is critical for tracking cells in aggregates and identifying co-occurring biomolecular events and factors that underlie the dynamics of the cytoskeleton, membrane lipids, second messengers, and gene expression. However, in *D. discoideum*, the number of frequently used FP species is limited to two or three. In this study, we explored the use of new-generation FP for practical 4- to 5-color fluorescence imaging of *D. discoideum*. We showed that the yellow fluorescent protein Achilles and the red fluorescent protein mScarlet-I both yield high signals and allow sensitive detection of rapid gene induction. The color palette was further expanded to include blue (mTagBFP2 and mTurquosie2), large Stoke-shift LSSmGFP, and near-infrared (miRFP670nano3) FPs, in addition to the HaloTag ligand SaraFluor 650T. Thus, we demonstrated the feasibility of deploying 4- and 5- color imaging of *D. discoideum* using conventional confocal microscopy.

## Introduction

Live-cell fluorescence imaging is indispensable for studying the functions of various biomolecules and organelles in cells and tissues. Genetically encoded fluorescent proteins (FP) expressed under specific promoters serve as surrogates to monitor gene expression or track intracellular localization as protein tags. Furthermore, FPs can be used as donor-acceptor pairs for Fluorescence Resonance Energy Transfer (FRET) to measure molecular interactions. Model systems in cell and developmental biology have incorporated new-generation FPs with superior stability and high quantum yields (Rodriguez *et al*., 2017; M. Wang *et al*., 2023) to encompass a wide range of visible light spectra from blue to near-infrared (NIR). In budding yeast, the nuclei, cell membranes, kinetochores, tubulins, and F-actin were simultaneously imaged using five FPs (Sakai *et al*., 2021): mTagBFP2 (Subach *et al*., 2011), mTurquoise2 (Goedhart *et al*., 2012), mNeonGreen (Shaner *et al*., 2013), mCherry (Shaner *et al*., 2004), and iRFP (Filonov *et al*., 2011). In *Neurospora crassa*, four FPs, mTagBFP2, mNeonGreen, mApple (Shaner *et al*., 2008), and iRFP670 (Shcherbakova and Verkhusha, 2013) were employed to visualize the nucleus, nuclear pore complex, microtubules, and cell polarity markers (Z. Wang *et al*., 2023). In *Caenorhabditis elegans*, four FPs, mTagBFP2, TagRFP-T (Shaner *et al*., 2008), mNeptune2.5 (Chu *et al*., 2014), and CyOFP1 (Chu *et al*., 2016) with a genetically encoded Ca^2+^ indicator GCaMP6s (Chen *et al*., 2013) were expressed simultaneously to identify individual neurons and monitor their activities (Yemini *et al*., 2021). These studies are only a small representation of works that show the benefit of utilizing a wide range of fluorescence spectra (i.e., the “color palette”) in imaging-based analysis.

*Dictyostelium discoideum* is a valuable model system for studying cytokinesis, phagocytosis, cell migration, and chemotaxis, along with other processes such as cell–cell signaling associated with its unique aggregative multicellularity. Although various FP has been utilized in *D. discoideum* (Table S1), fluorescence imaging using three or more fluorescent markers has not been widely adopted. mTurquoise2, the bright variant of CFP fused to cAMP receptor cAR1, expressed together with RegA-mRFPmars and AcaA-GFP highlighted cell–cell heterogeneity at the early gene expression level (Mukai *et al*., 2016). The CFP- and YFP-based FRET measurements of cytosolic cAMP combined with the detection of RFP-tagged MS2 coat protein (Masaki *et al*., 2013) or PH domain fused to RFP to monitor cAMP-induced synthesis of mRNA and phosphatidyl inositol (3,4,5) trisphosphate (PIP3) (Kamino *et al*., 2017) characterized the coupling between cell–cell signaling, gene expression, and cell migration. The FP repertoire has since expanded to include short wavelength blue fluorescent TagBFP (Kundert *et al*., 2020) and NIR fluorescent protein iRFP (Ohta *et al*., 2018); however, these two ends of the spectrum have not been utilized in *Dictyostelium* except in a case of 4-color imaging (Kundert *et al*., 2020) with mCerulean (Rizzo and Piston, 2005), Superfolder GFP (Pédelacq *et al*., 2005), Flamindo2 (Odaka *et al*., 2014), and mCherry.

In *D. discoideum*, one of the obstacles to live-cell imaging is its high photosensitivity to blue light (Zhang *et al*., 2023), which precludes the repeated application of strong excitation light. Earlier generations of FPs have relatively low quantum yields (Matlashov *et al*., 2020; Seo *et al*., 2024) making long-duration time-lapse imaging unfeasible. Here, we tested newer-generation FPs, either alone or in a tagged format, to increase the practicality of 4- and 5-color fluorescence imaging in *Dictyostelium*. We examined mTagBFP2, Achilles (Yoshioka-Kobayashi *et al*., 2020), mScarlet-I (Bindels *et al*., 2016) and miRFP670nano3 (Oliinyk *et al*., 2022), for brightness and intracellular localization in tagged form, and compared them with conventionally used FPs. By employing these FPs along with the large Stoke shift LSSmGFP (Campbell *et al*., 2022), we demonstrated practical 4- and 5-color fluorescence live-cell confocal imaging in *D. discoideum*.

## Materials & Methods

### Plasmids and transformation

Plasmids constructed in this study are summarized in Table S2. For constitutive expression, the majority of vectors employed *act15* promoter, except some vectors that employed *coaA* promoter (Table S2). For details, see Supplementary Information. To introduce extrachromosomal vectors, Ax4 cells were electroporated with 1 µg plasmid DNA following the standard protocol (Nellen *et al*., 1984). Transformants were selected in HL5 growth medium containing 10 µg/mL G418, 60 µg/mL Hygromycin B and 10LJµg/mL Blasticidin S either alone or in combination. For 4- and 5- color imaging, we introduced single vectors (G418 resistance) carrying two tags along with the knock-in of mScarlet-I-tagged PKBR1(N150) to *act5* locus (see Supplementary Information).

### Cell culture, development, and labeling

Cells were grown in HL5 medium at 22 LJ either in shake flasks or in petri dishes. The growth media contained 10 µg/mL G418, 60 µg/mL Hygromycin B, and 10 µg/mL Blasticidin S where appropriate. To image the growth stage, the cells were collected, washed twice, and resuspended in Phosphate Buffer (PB: 12 mM KH_2_PO_4_, 8 mM Na_2_HPO_4_, pH 6.5). For NIR imaging, cells expressing miRFP670nano3 and HaloTag were labeled with biliverdin and NIR HaloTag ligands, respectively (Supplementary Information). A drop of cell suspension was plated either directly onto a 24×50 mm coverslip (Matsunami) or a φ25 mm round coverslip (Matsunami) mounted on a metal chamber (Attofluor, Invitrogen). The cells were allowed to attach to the substrate before image acquisition. To image slug and fruiting bodies, washed cells were suspended at the density of 2×10^7^ cells/mL in PB, and 5 µL of the cell suspension was deposited on an agar plate (2% agar (Bacto) in Milli-Q water). For cellulose labeling, the cells were developed on an agar plate containing a cellulose-staining dye (see Supplementary Information). After 15–24 h, samples were excised together with the agar sheet and placed upside down onto a coverslip with a 50 µm height polyethylene terephthalate spacer ring (vinyl patch transparent Ta-3N, Kokuyo). The inner space of the ring was filled with liquid paraffin (Nacalai Tesque, Light 26132) (Hashimura *et al*., 2019) before observation.

The following procedure was followed to observe cells dissociated from the slugs. Growing cells were collected and suspended in PB at a density of 10^7^ cells/mL. Then, 1.5 mL of cell suspension was deposited on a 60 mm agar plate (2% agar (Bacto) in Milli-Q water) containing 25 µg/mL biliverdin. Excess water was removed after the cells attached to the substrate. After 20 h of incubation at 22 °C, slugs were collected and passed through a 23G syringe needle (NN-2332R, Terumo) 10 times; approximately 0.4 mL of cell suspension at the density of 1.5 ×10^5^ cells/mL was loaded into a custom-made microfluidic chamber with a 2.5 µm height observation channel (Fujimori *et al*., 2019). The cells were allowed to attach for 30 min prior to observation.

### Image acquisition and analysis

To observe the cells expressing the two FPs, green and red (Fig. 1, 2, 3A, 4, 5, 6, S1 and S2), an inverted microscope (Ti-E, Nikon) equipped with a laser confocal point-scanning unit (A1R, Nikon) was employed. A 488 and 561 nm laser was used for excitation, together with a multiband 405/488/561 nm dichroic mirror and 525/50 and 595/50 bandpass filters for detection. For 3D time-lapse imaging in Fig. 5, 13 z-section images were taken at 7-µm-intervals with a piezo stage (Nano-Drive, Mad City Labs). For observations that included blue and/or near-IR (Fig. 7, 9, 10A, 11, S3, and S4), a multibeam confocal scanning unit (CSU-W1, Yokogawa) and an electron-magnified CCD camera (iXon 888, Andor) on an inverted microscope (IX83, Olympus) were employed. 405, 445, 515, 560, and 638 nm lasers were used for excitation, together with a Di01-T405/488/568/647 dichroic beam splitter (Semrock) and appropriate emission filters. To observe the cells expressing LSSmGFP (Fig. 8 and Fig 10B), a multibeam confocal scanning unit (CSU-W1, Yokogawa) and CMOS cameras (ORCA-Fusion BT, Hamamatsu) on an inverted microscope (IX83, Olympus) were employed. To check the LSSmGFP fluorescence (Fig. 8), 405 and 488 nm lasers were employed together with a multiband dichroic mirror (T405/488/561/640 nm) and 447/60 and 525/50 bandpass filters. For 5-color imaging (Fig 10B), 405, 488, 561 and 640 nm lasers were used for excitation, and 447/60, 525/50, 617/73, and 685/40 bandpass filters were used for detection.

**Fig. 1.**
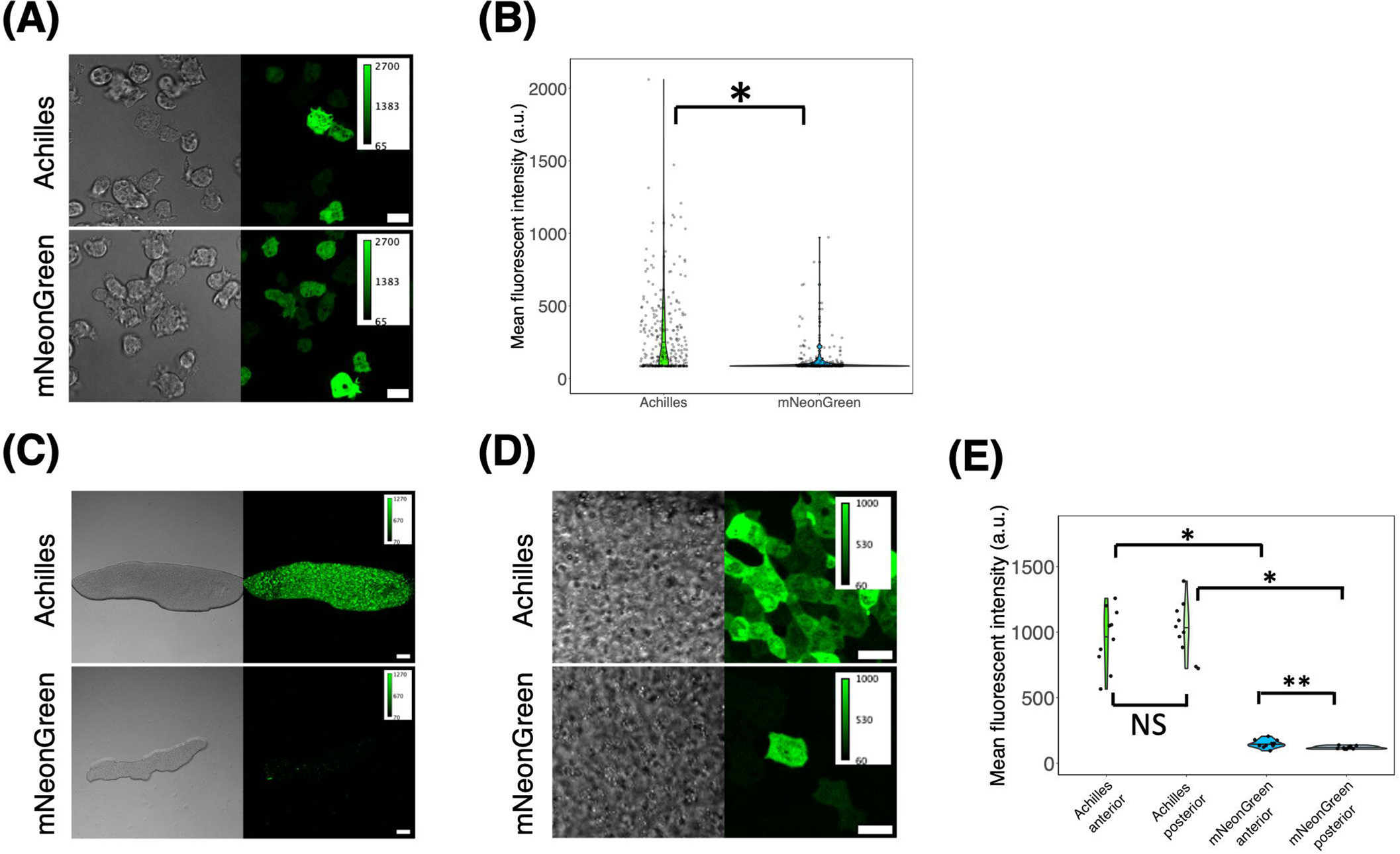
Achilles as a superior green fluorescent protein alternative in D. discoideum. (A) Vegetative cells expressing Achilles (upper panels) and mNeonGreen (lower panels) under *act15* promoter (left: brightfield channel, right: green channel). Scale bar, 10 µm. (B) Violin plot of the single-cell mean fluorescence intensity distribution of Achilles and mNeonGreen expressing vegetative cells (Achilles, n = 297 cells. mNeonGreen, n = 266 cells). The black line is the median. *: *P* < 10 ^-15^. (C) A slug harboring Achilles (upper panels) and mNeonGreen (lower panel) *act15* promoter expression plasmids (left: brightfield channel, right: green channel). Scale bar, 100 µm. (D) High magnification images of slugs harboring Achilles (upper panels) and mNeonGreen (lower panels) *act15* promoter expression plasmids (left: brightfield channel, right: green channel). The anterior-posterior axis of the slug is from left to right. Scale bar, 10 µm. (E) Violin plot of fluorescent intensities measured in the anterior and the posterior region of a slug (Achilles, n = 10 slugs. mNeonGreen, n = 10 slugs). The black line indicates the median value.*: *P* < 10 ^-4^. **: *P* < 0.05. NS: not significant (*P* > 0.05).

**Fig. 2.**
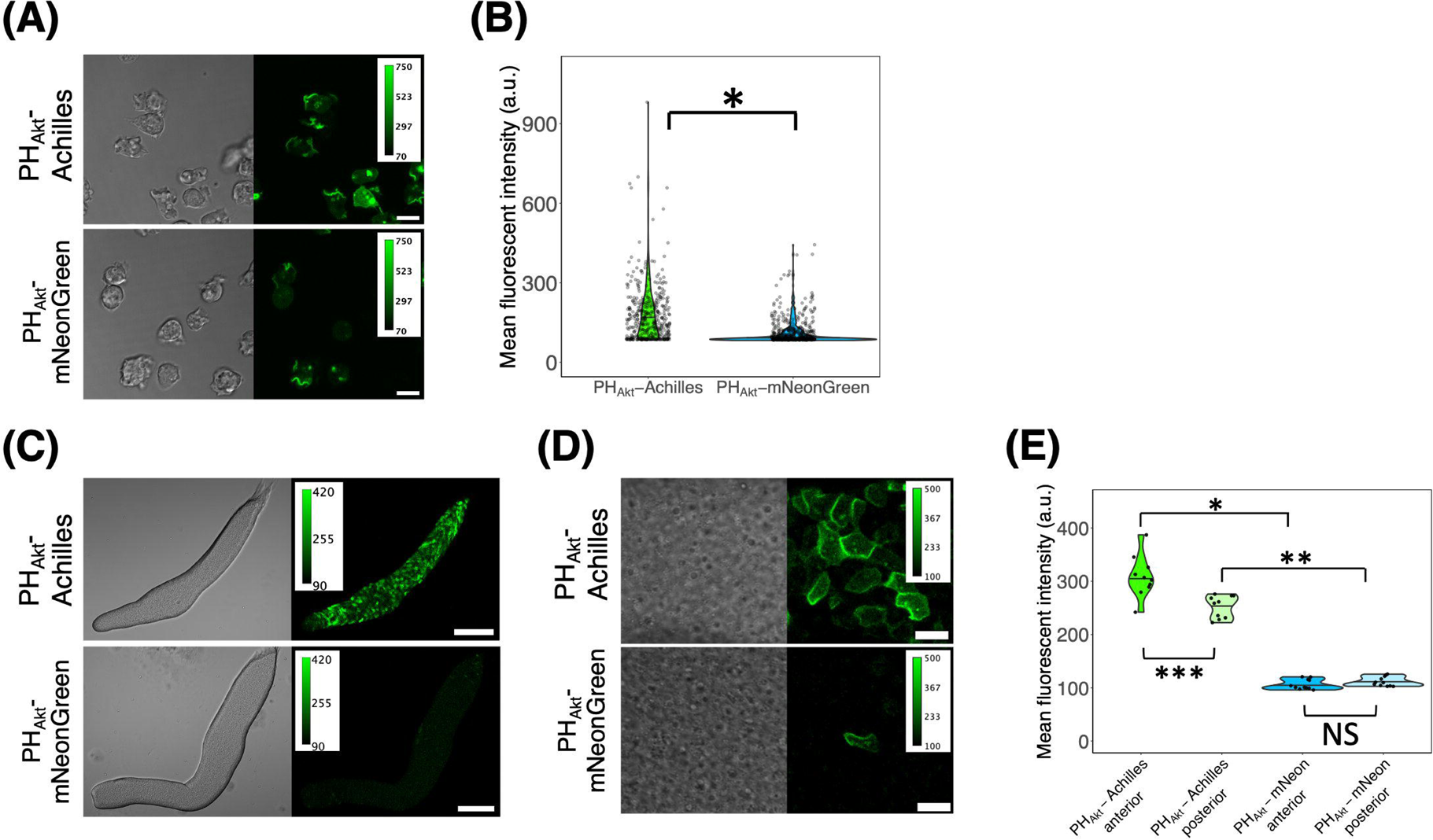
Use of Achilles as a yellow fluorescent protein tag in *D. discoideum*. (A) Cells carrying *coaAp*:PH_Akt_-Achilles (upper panels) and *coaAp*:PH_Akt_-mNeonGreen (lower panels) (left: brightfield channel, right: green channel). Scale bar, 10 µm. (B) Violin plot of the single-cell mean fluorescence intensity distribution of PH_Akt_-Achilles and PH_Akt_-mNeonGreen expressing vegetative cells (PH_Akt_-Achilles, n = 402 cells. PH_Akt_-mNeonGreen, n = 599 cells). The black line indicates the median. *: *P* < 10 ^-15^. (C) Slugs (PH_Akt_-Achilles or -mNeonGreen under *coaA* promoter. Scale bar, 100 µm. (D) High magnification images of a slug (upper panel PH_Akt_-Achilles, lower panels PH_Akt_-mNeonGreen). The anterior-posterior axis of the slug is from left to right. Scale bar, 10 µm. (E) Violin plot of the mean fluorescent intensities of the anterior and the posterior region of a slug (PH_Akt_-Achilles, n = 12 slugs. PH_Akt_-mNeonGreen, n = 12 slugs). The black line indicates the median value.*: *P* < 10 ^-5^. **: *P* < 0.05. ***: *P* < 0.01. NS: not significant (*P* > 0.05).

**Fig. 3.**
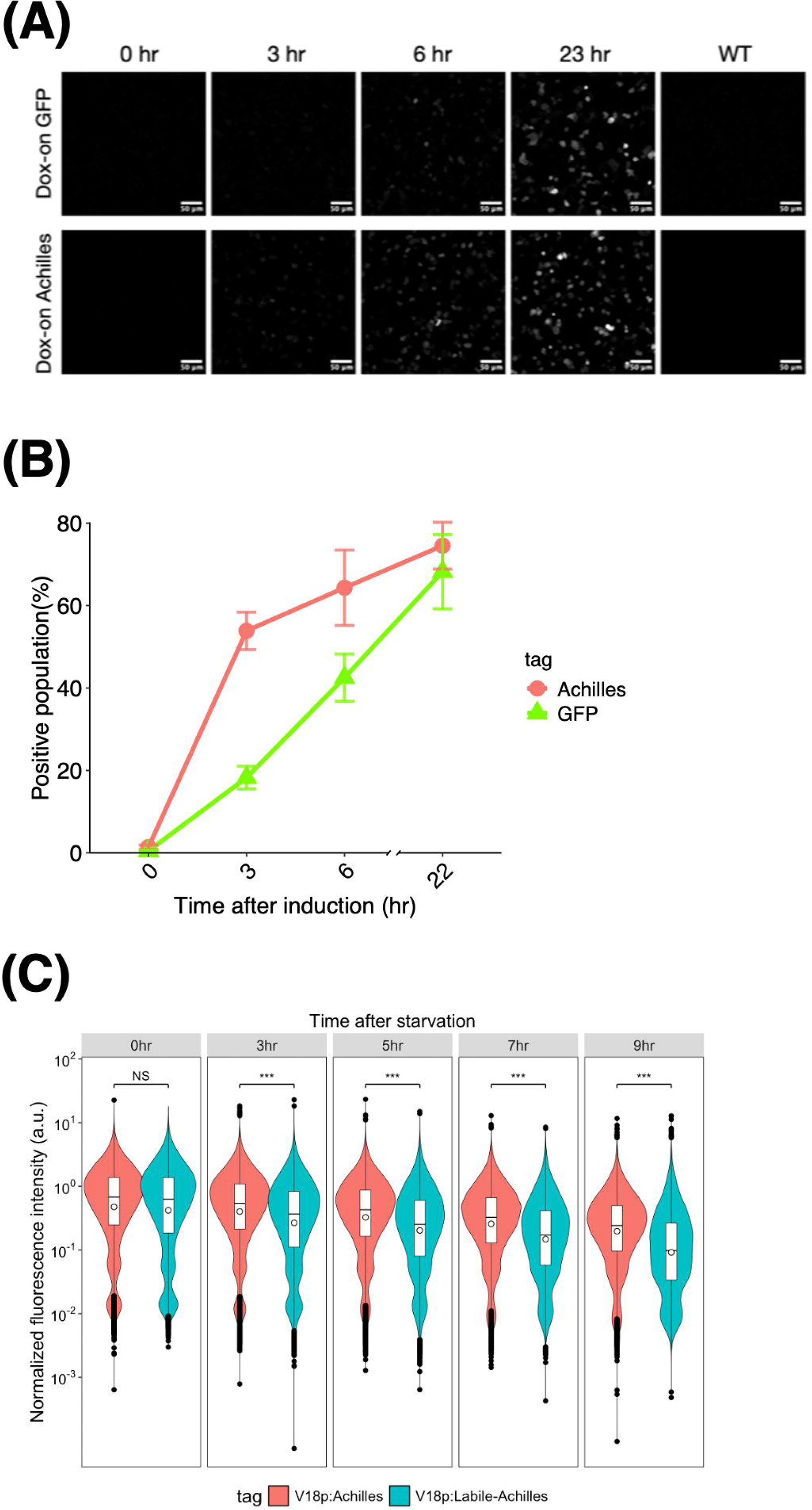
Achilles and labile-Achilles as fast responding reporter genes in *D. discoideum*. (A) Dox-induced GFP(S65T) (upper panel) and Achilles (lower panel) fluorescence in vegetative cells plated from shaken culture. Cells were incubated with 10 µg/mL Dox for indicated time (0, 3, 6, and 22 h). WT indicates parental Ax4 cells without expression of fluorescence protein. (B) Percentages of the cells displaying GFP(S65T) or Achilles fluorescence after doxycycline (Dox) addition. Data obtained through flow cytometry (see Material and Methods). Dots and error bars represent the average of positive cells and their standard deviation (three independent experimental runs; n = 135,000 cells, 45,000 cells per experiment). (C) Flow cytometry of *V18p*:Achilles and *V18p*:Labile-Achilles. The violin plots represent three independent experimental runs, totaling 90,000 cells (3,000 cells per experiment). The boxes and whiskers indicate the mean values and the interquartile ranges, respectively. Outliers are represented as dots. A Welch Two Sample t-test was conducted to compare the two groups, and significance levels are indicated (***: *P* < 2.2e-16).

**Fig. 4.**
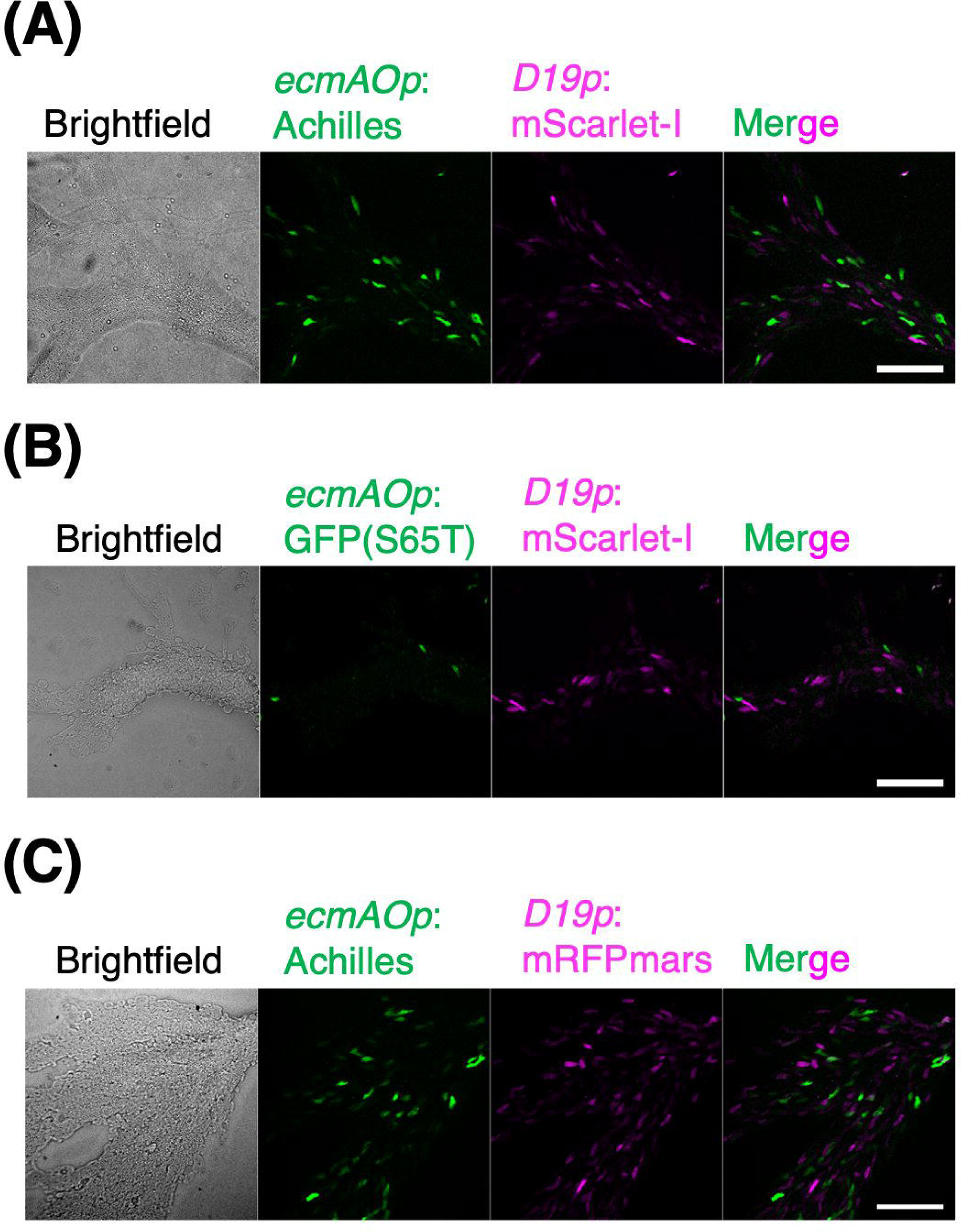
Yellow fluorescent protein Achilles allows early detection of developmental gene expression *D. discoideum*. (A–C) Streaming-stage cells (8 h after starvation) co-expressing green and red FPs under prestalk and prespore specific-promoter. Panels from left to right: Brightfield images, green fluorescence, red fluorescence and merged image. Scale bars, 100 µm. (A) *ecmAOp*:Achilles (green) and *D19p*:mScarlet-I (magenta). (B) *ecmAOp*:GFP(S65T) (green) and *D19p*:mScarlet-I (magenta). (C) *ecmAOp*:Achilles (green) and *D19p*:mRFPmars (magenta).

**Fig. 5.**
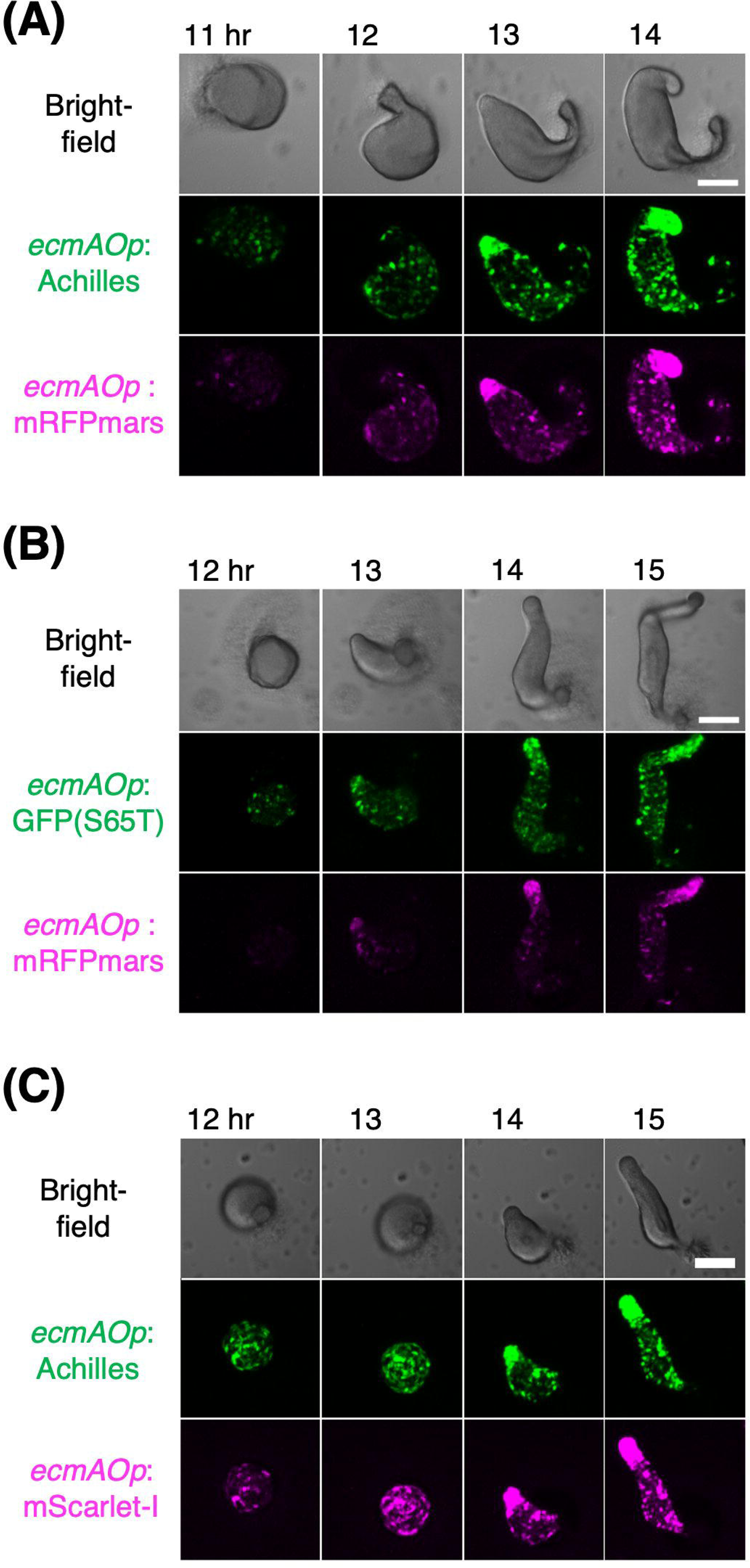
Red fluorescent protein mScarlet-I allows early detection of developmental gene expression in *D. discoideum*. (A–C) Cells co-expressing green and red FPs under prestalk-specific promoter. The numbers indicate hours after nutrient removal. Brightfield images (top panels). Maximum intensity projection of z-stacks (middle and bottom panels). Scale bar, 100 µm. (A) *ecmAOp*:Achilles (green) and *ecmAOp*:mRFPmars (magenta). (B) *ecmAOp*:GFP(S65T) (green) and *ecmAOp*:mRFPmars (magenta). (C) *ecmAOp*:Achilles (green) and *ecmAOp*:mScarlet-I (magenta).

**Fig. 6.**
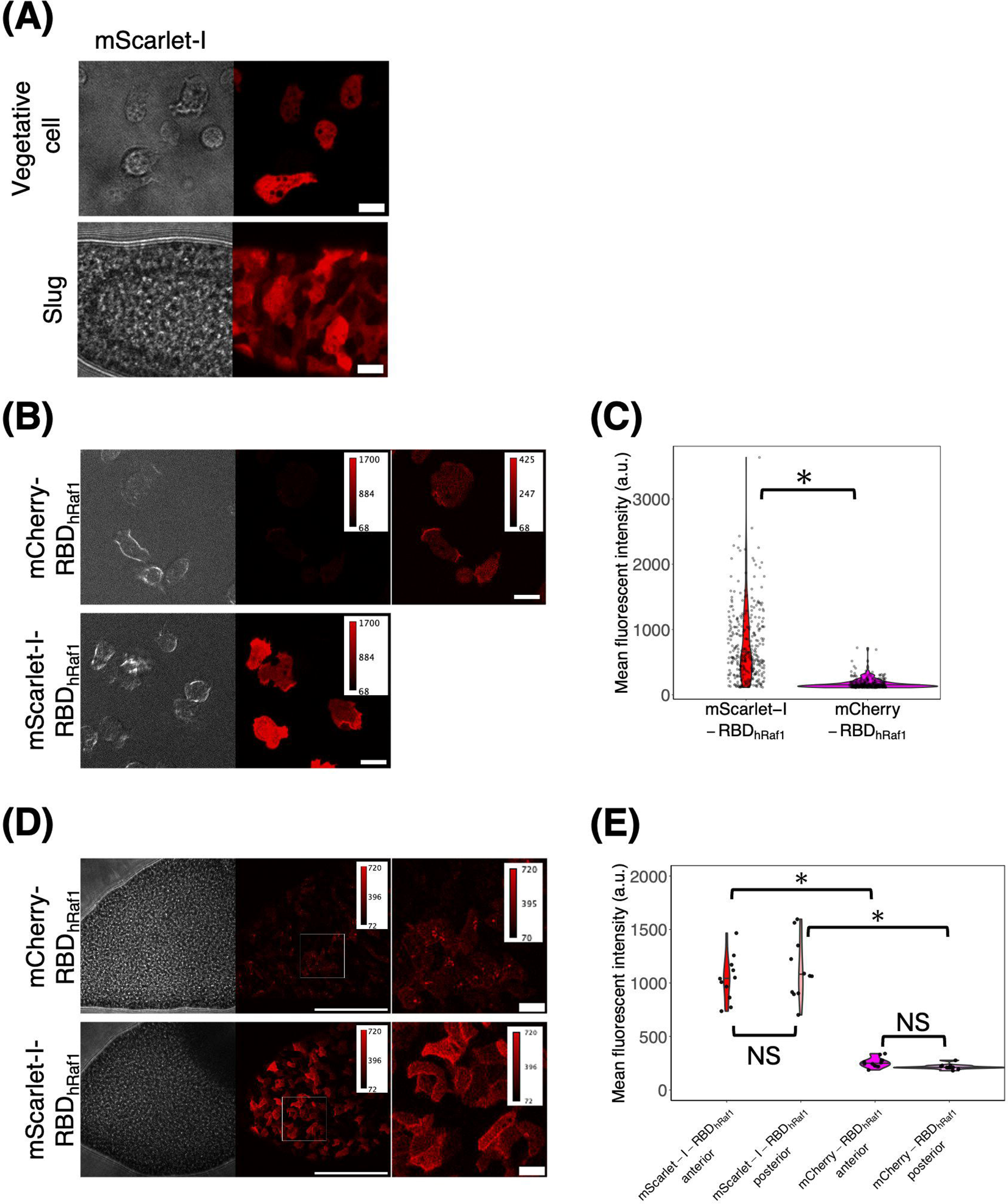
mScarlet-I is a first-choice red fluorescent protein for protein tagging in *D. discoideum*. (A) *act15p*:mScarlet-I cells. Vegetative cells (left panels) and cells in a slug (right panels). Brightfield (left panels) and fluorescence images (right panels). Scale bar, 10 µm. (B) *act15p*:mCherry-RBD_hRaf1_ (upper panels) and *act15p*:mScarlet-I-RBD_hRaf1_ (lower panel) expressing vegetative cells. Contrast was adjusted as indicated (top right; color bars). Scale bar, 10 µm. (C) Violin plots of the mean fluorescence intensity of mCherry-RBD_hRaf1_ and mScarlet-I-RBD_hRaf1_ expressing vegetative cells (mCherry-RBD_hRaf1_, n = 298 cells. mScarlet-I-RBD_hRaf1_, n = 333 cells). Black line indicates the median. *: *P* < 10 ^-15^. (D) The slug anterior-region of *act15p*:mCherry-RBD_hRaf1_ and *act15p*:mScarlet-I-RBD_hRaf1_ cells. Magnified view (right panels) of the white-boxed area (left panels). The anterior-posterior axis of the slug is from left to right. Scale bar, 100 µm (left) and 10 µm (right). (E) Violin plots of fluorescence intensity distribution in the anterior and the posterior region of the slug expressing mCherry-RBD_hRaf1_ or mScarlet-I-RBD_hRaf1_ (mCherry-RBD_hRaf1_, n = 11 slugs. mScarlet-I-RBD_hRaf1_, n = 12 slugs). The black line indicates the median. *: *P* < 10 ^-5^. NS: not significant (*P* > 0.05).

**Fig. 7.**
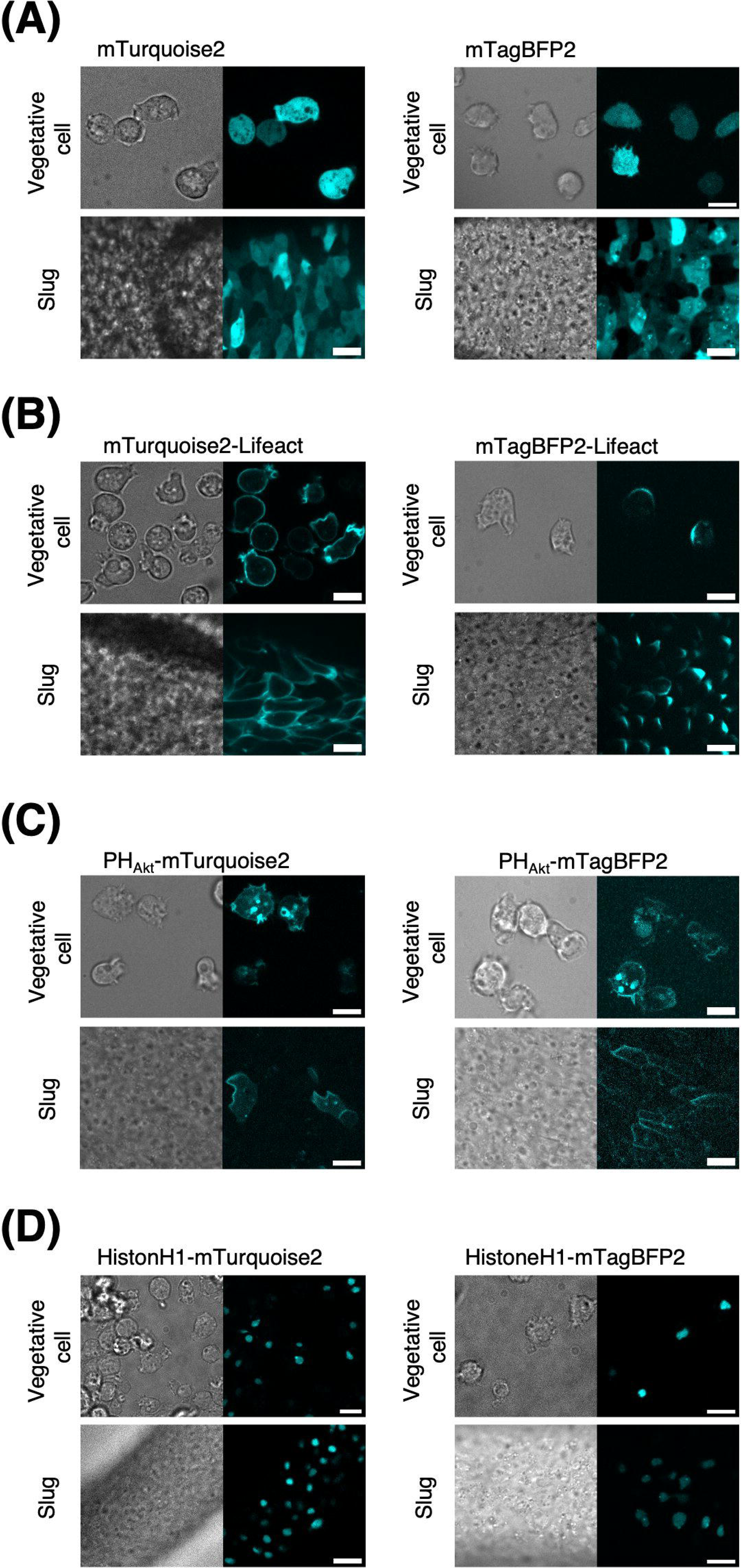
Blue fluorescent proteins mTurquoise2 and mTagBFP2 for live-cell imaging in *D. discoideum*. (A–H) The brightfield (left panels) and fluorescence images (right panels) of blue fluorescent protein expressing vegetative cells (upper panels) and cells in a slug (lower panels). The anterior-posterior axis of the slug is from left to right. Scale bar, 10 µm. (A) *act15p*:mTurquoise2 and (B) *act15p:*mTagBFP2 vegetative cells. (C) *act15p:*mTurquoise2-Lifeact and (D) *act15p:*mTagBFP2-Lifeact. (E) *coaAp*:PH_Akt_-mTurquoise2 and (F) *coaAp*:PH_Akt-_mTagBFP2. (G) *act15p*:HistoneH1-mTurquoise2 and (H) *act15p*:HistoneH1-mTagBFP2.

**Fig. 8.**
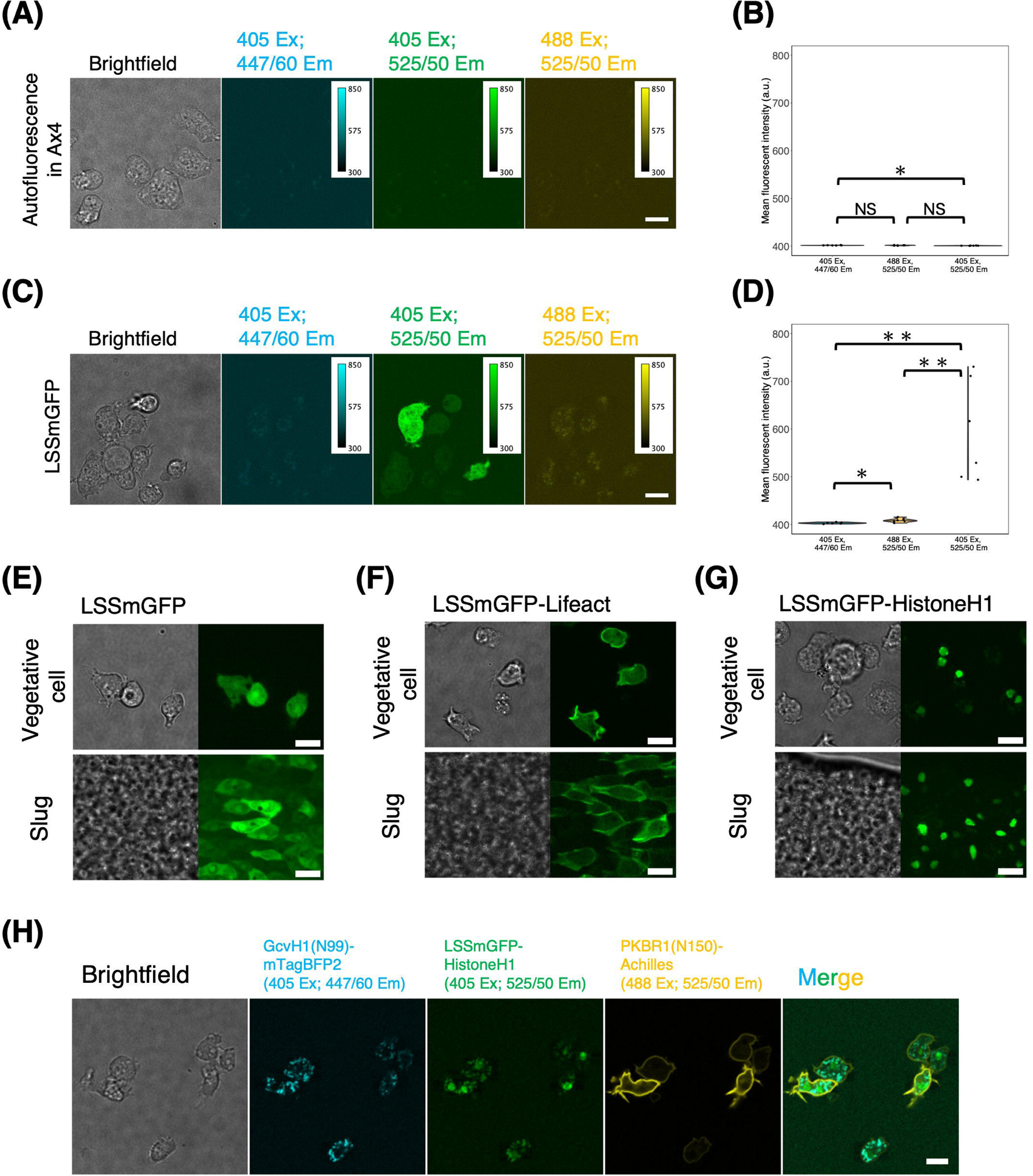
Live-cell imaging using large stokes shift LSSmGFP in *D. discoideum*. (A, B) *act15p*:LSSmGFP and (C, D) control Ax4 cells in the vegetative stage. (A, C) From left to right; brightfield, fluorescent images (405 nm excitation, 447/60 nm emission), (405 nm excitation, 525/50 nm emission), (488 nm excitation, 525/50 nm emission). Scale bars, 10 µm. (B, D) Violin plots of the mean fluorescence intensity of Ax4 (B) or LSSmGFP expressing vegetative cells (D) (Ax4, n = 6 cells. LSSmGFP, n = 6 cells). Black line indicates the median. *: *P* < 0.05. **: *P* < 0.01. NS: not significant (*P* > 0.05). (E–G) Fluorescent images of *act15p*:LSSmGFP (E), *act15p*:LSSmGFP-Lifeact (F) and *act15p*:LSSmGFP-HistoneH1 (G). Vegetative cells (top) and the slug cells (bottom). Brightfield (left panels) and fluorescent images (405 nm excitation, 525/50 nm emission; right panels). The anterior-posterior axis of the slug is left to right. Scale bar, 10 µm. (H) 3-color fluorescence images. (Left to right panels) Brightfield, mitochondria marker *act15p*:GcvH1(N99)-mTagBFP2, nucleus *act15p*:LSSmGFP-HistoneH1, plasma membrane *coaAp*:PKBR1(N150)-Achilles and merged fluorescent channels. Scale bar, 10 µm.

**Fig. 9.**
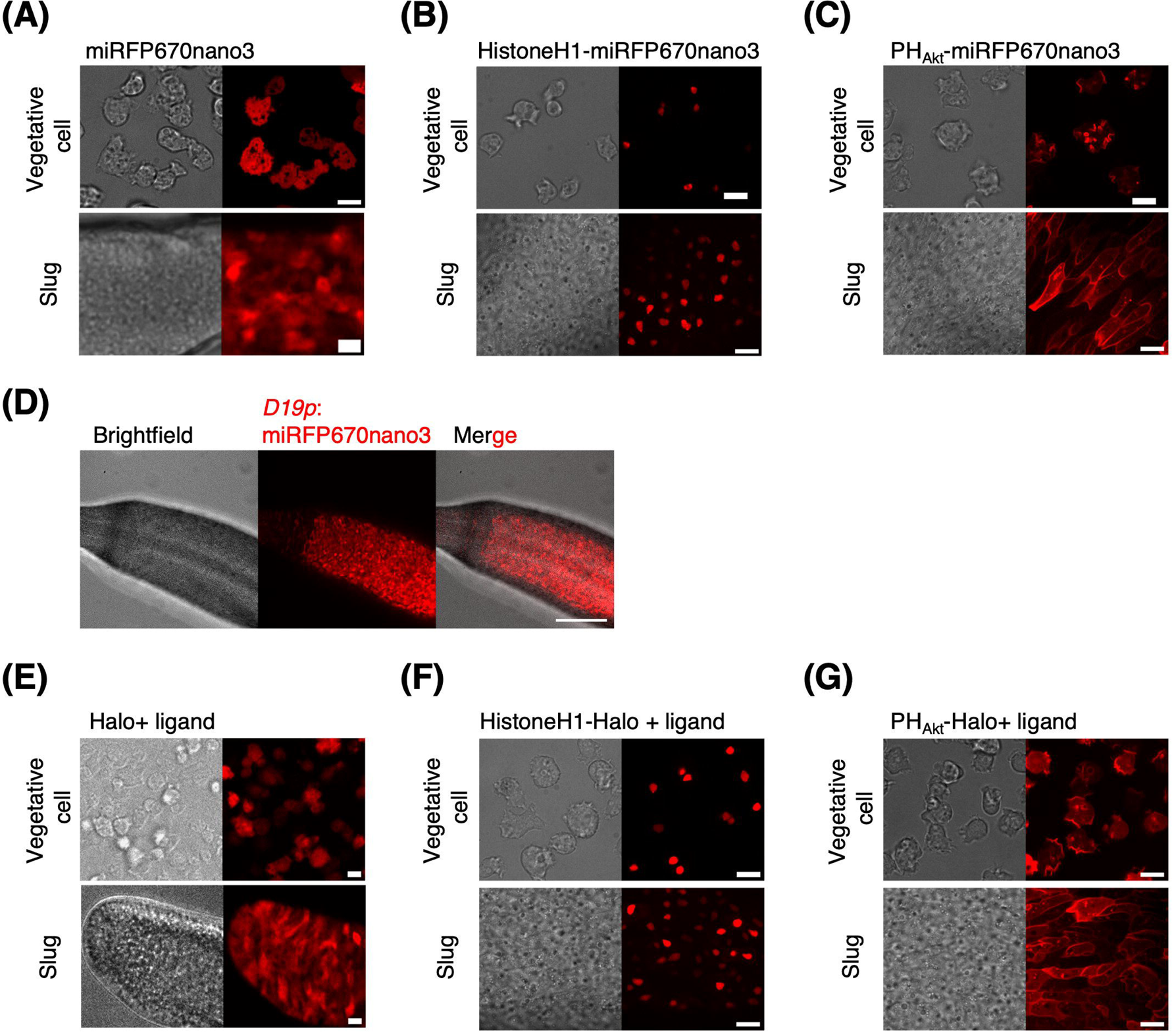
Live-cell imaging of *D. discoideum* in near-IR spectrum. (A–C) Snapshots of vegetative cells expressing *act15p*:miRFP670nano3 (A), *act15p*:HistoneH1-miRFP670nano3 (B) and *coaA*:PH_Akt_-miRFP670nano3 (C). Brightfield (left panels) and fluorescence images (right panels). Scale bars, 10 µm. (D) A slug expressing miRFP670nano3 under the prespore-specific *D19* promoter. Brightfield (left), near-IR fluorescence image (middle), merged image (right). Scale bar, 100 µm. (E–G) Snapshots of vegetative cells expressing *act15p*:Halo (E), *act15p*:HistoneH1-Halo (F) and *coaA*:PH_Akt_-Halo (G). Left panels, bright field. Right panels near-IR fluorescence of Halo-ligand SaraFluor650T. Scale bars, 10 µm. The anterior-posterior axis of the slug is from left to right (A-G).

**Fig. 10.**
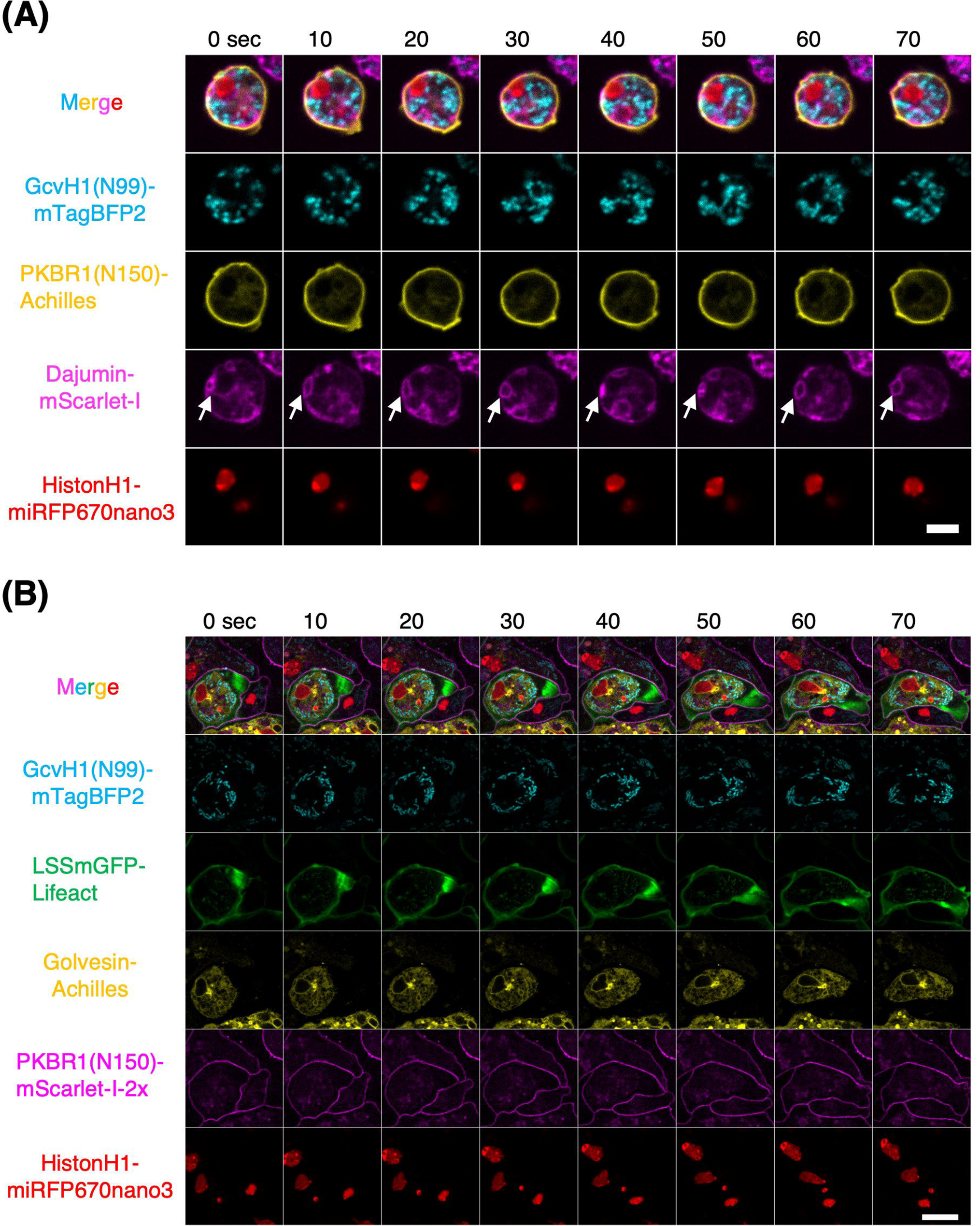
4- and 5-color fluorescence live-cell imaging in *D. discoideum*. (A) Vegetative cells expressing 4 FP-tags. From top to bottom: merged channel, *act15p*:GcvH1(N99)-mTagBFP2 (mitochondria; cyan), *coaAp*:PKBR1(N150)-Achilles (plasma membrane; yellow), *act15p*:Dajumin-mScarlet-I (contractile vacuole; magenta), and *act15p*:HistoneH1-miRFP670nano3 (nuclei; red). White allows: contractile vacuole going through cycles of deformation. Scale bar, 5 µm. (B) Slug cells expressing 5-FP-tags. From top to bottom: merged channel, *act15p*:GcvH1(N99)-mTagBFP2 (mitochondria; cyan), *act15p*:LSSmGFP-Lifeact (F-actin; green), *act15p*:Golvesin-Achilles (Golgi apparatus and ER network; yellow), *act5p*:PKBR1(N150)-mScarlet-I-2x (plasma membrane; magenta) and *act15p*:HistoneH1-miRFP670nano3 (nucleus; red). Scale bar, 10 µm.

**Fig. 11.**
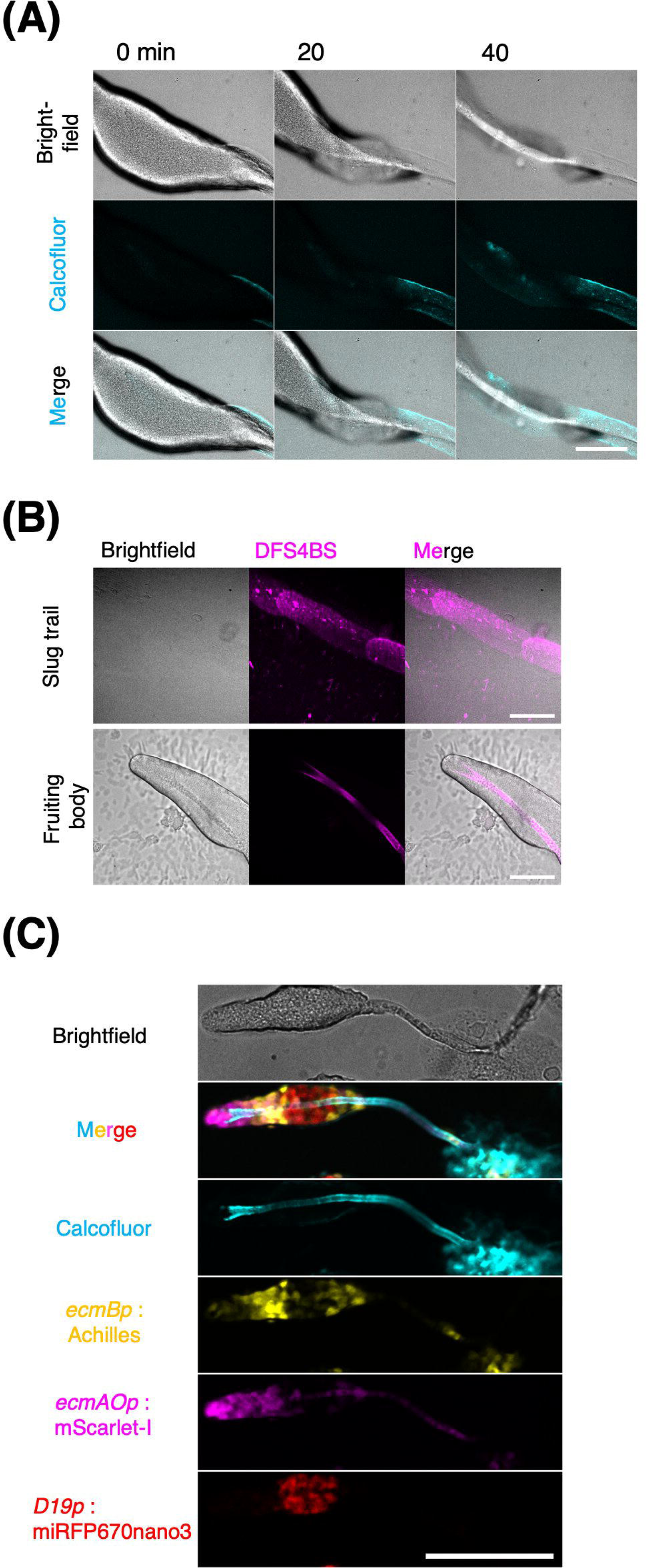
Combining cellulose staining with FP-tags to image multicellular body of *D. discoideum*. (A) A migrating slug stained with cellulose-specific dye calcofluor white. Brightfield (top), fluorescence (middle), and merged images (bottom). The slug migrated toward the left side of the images. Scale bar, 100 µm. (B) Time-lapse imaging of culminating *D. discoideum* stained with cellulose-specific dye Direct Fast Scarlet 4BS (DFS4BS). Brightfield (left), DFS4BS (middle), and merged (right). Slug stage (upper panels) with cellulose-rich trailing slime sheath on agar. A culminant (lower panels) with a stalk tube. Scale bar, 100 µm. (C) Culminating *D. discoideum* stained with calcofluor white (cyan) expressing Achilles (PstB cell; yellow), mScarlet-I (PstA and PstO cell; magenta), and miRFP670nano3 (prespore cell; red) under *ecmB, ecmAO,* and *D19* promoter, respectively. Scale bar, 100 µm.

Images were analyzed using ImageJ and R statistical packages. To quantify the mean fluorescence intensity of individual cells, cell masks were obtained from brightfield images using Trainable Weka Segmentation (Arganda-Carreras *et al*., 2017). To quantify fluorescence of slugs, the mean fluorescence intensity of 30 µm^2^ rectangular region at the anterior or posterior region of the slug was measured. For 5-color fluorescence imaging, bleed-through of LSSmGFP was corrected (see Supplementary information). In all statistical analyses, *P* values were determined using the Wilcoxon rank-sum test with Bonferroni adjustment.

### Measurement of maturation and turn-over of EGFP and Achilles

For the induction experiments (Fig. 3A and B), Ax4 cells were transformed with either a doxycycline (Dox)-inducible GFP expression vector, pDM340 (Veltman, Keizer-Gunnink, *et al*., 2009) or a doxycycline-inducible Achilles expression vector (Supplementary information); 10 µg/mL (final conc.) doxycycline was added to the shaken cultures and incubated for 0, 3, 6, or 22 h. Cells were then collected and washed twice with PB, resuspended in 0.3 mM EDTA containing PB, and the fluorescence intensity was measured using a flow cytometer (Sony LE-SH800AC; excitation 488 nm, 525/50 nm bandpass filter). Gating was set based on both forward scatter (FSC) and backward scatter (BSC). Cells with a fluorescence intensity three times the standard deviation above the mean (mean + 3×SD) of the 0 h cells were counted as positive. The fluorescence signal at each time point was divided by the intensity at 0 h for normalization.

For labile-Achilles fluorescence quantification (Fig. 3C), cells expressing either Achilles or labile-Achilles under *V18* promoter were collected at mid-log phase, washed twice by centrifugation, and resuspended in PB. For cytometry of vegetative cells, washed cells were suspended in PB containing 0.3 mM EDTA for loading. For cytometry of developing cells, washed cells (2 × 10^7^ cells/mL in PB) were plated onto a nitrocellulose filter on top of an absorbent pad soaked in PB and incubated in the dark at 22 °C. After 3, 5, 7, and 9 h, the cells were harvested by gentle scraping using a syringe needle and suspended in 20 mM EDTA containing PB in 1.5 mL tubes. Cell aggregates were passed through a 23G syringe needle (Terumo) approximately 20 times before flow cytometry. Data processing and statistical analyses were performed using R statistical package.

## Results

### Expression of fast maturing bright FP Achilles and mScarlet-I in *D. discoideum*

Green fluorescent proteins utilized in *D. discoideum* today are mostly spectral variants of the GFP of *Aequorea victoria* (Chalfie *et al*., 1994; Kataria *et al*., 2013; Mukai *et al*., 2016) selected for reduced crossover of excitation and emission spectra (Goedhart *et al*., 2012; Nagai *et al*., 2002). Additionally, mNeonGreen derived from *Branchiostoma lanceolatum* which is three times brighter than GFP (Shaner *et al*., 2013) has been adopted more recently (Honda *et al*., 2021; Nichols *et al*., 2020; Paschke *et al*., 2018). To update the FP repertoire in *D. discoideum*, we first studied the stability and expression of Achilles, a yellow fluorescent protein derived from Venus, for fast maturation (Yoshioka-Kobayashi *et al*., 2020). We introduced an extrachromosomal vector that drives Achilles or mNeonGreen expression under the control of the strong *act15* promoter. Using a confocal microscope (see Materials & Methods), the fluorescence of both Achilles and mNeonGreen appeared uniformly in the cytosol and nucleus of vegetative cells (Fig. 1A). The cells expressing Achilles appeared significantly brighter than those expressing mNeonGreen (Fig. 1B). In the slug stage, Achilles fluorescence remained bright in the majority of cells and appeared uniform in the cytosol (Fig. 1C and D), whereas very few cells showed mNeonGreen fluorescence (Fig. 1C–E).

Next, we compared Achilles and mNeonGreen protein tags. We first examined cells carrying an extrachromosomal expression vector for Achilles or mNeonGreen fused to the PH domain of Akt/PKB (Meili *et al*., 1999). The strong *coaA* promoter (Katoh-Kurasawa *et al*., 2021) was used for more homogeneous expression than in the *act15* promoter in the slug (Fig. S1). Using the same microscopy setup as above, in vegetative cells, PH_Akt_-Achilles and PH_Akt_-mNeonGreen fluorescence appeared localized in the pinocytic cups (Fig. 2A), as expected from an earlier observation of PH_Akt_-GFP (Ruchira *et al*., 2004). Cells expressing PH_Akt_-Achilles appeared brighter than those expressing PH_Akt_-mNeonGreen (Fig. 2B). In the slug stage, PH_Akt_-Achilles fluorescence was bright with small cell–cell variability, showing enrichment in the front cell–cell contact region (Fig. 2C and D). There was no visible effect on the ability of cells to form slugs or fruiting bodies. In contrast, PH_Akt_-mNeonGreen fluorescence was almost undetectable, except very rare cells that showed localized patterns similar to PH_Akt_-Achilles (Fig. 2C–E). This indicates that while Achilles behaves well during development, mNeonGreen may be less stable. Similarly, while both the N- and C-terminal fusions of Lifeact-mNeonGreen exhibited the expected localized patterns in vegetative cells (Fig. S2A and B), the fluorescence was markedly diminished in the slug stage of C-terminal fusion (Fig. S2C–E), suggesting that the free C-terminus of mNeonGreen may be subjected to regulated degradation in *D. discoideum*. These results indicate that, when studying later developmental stages, Achilles is the green/yellow FP of choice for brightness and stability.

Next, we studied whether the fast maturation of Achilles (Yoshioka-Kobayashi *et al*., 2020) could be used in *D. discoideum*. To measure the rate of increase in fluorescence intensity, strains carrying a dox-on-inducible vector (Veltman, Keizer-Gunnink, *et al*., 2009) with either GFP or Achilles under an inducible promoter were employed. Confocal microscopy images showed a clear increase in fluorescence after induction with doxycycline in both the strains (Fig. 3A). Flow cytometry was performed for quantification (see Materials and Methods). While 64% of the Dox-Achilles cells took only 6 h to show detectable fluorescence (Fig. 3B, red), 22 h was required for a similar percentage of Dox-GFP(S65T) cells (68%) to display a comparable level of fluorescence intensity (Fig. 3B, green). These result suggests that despite the low temperature (22 °C) of *D. discoideum* culture, maturation of Achilles was fast and comparable to that demonstrated in mice (Yoshioka-Kobayashi *et al*., 2020). Rapid maturation makes Achilles ideal for the sensitive detection of early gene expression in *D. discoideum*. To further extend its applications, we sought to reduce its half-life for a rapid response to gene downregulation (Sunabori *et al*., 2008). Based on labile-GFP (Deichsel *et al*., 1999), we constructed “labile-Achilles” where the N-terminus of Achilles was tagged to the first 33 amino acids of *S. cerevisiae* ubiquitin (Bachmair *et al*., 1986). We tested its half-life by expressing labile-Achilles under *V18* promoter, whose activity diminishes after starvation (Singleton *et al*., 1989). Fig. 3C shows a plot of the intensity of Achilles and labile-Achilles-expressing cells developed on a nitrocellulose filter (see Materials and Methods). A difference in the reduction of Achilles and labile-Achilles fluorescence was detectable during the first 3 h after starvation. After 5 h, the fluorescence of labile-Achilles decreased by 46% (Fig. 3C, 5 h), whereas it took approximately 7 h for Achilles to reach a comparable level (Fig. 3C, 7 h). These results suggest that labile-Achilles is a fast-responding alternative to labile-GFP for monitoring promoter activity.

Next, we explored the use of Achilles and red FP mScarlet-I as cell-type markers. mScarlet-I is a fast-maturing variant of mScarlet that was developed from synthetic constructs to yield high brightness while avoiding dimerization (Bindels *et al*., 2016). We constructed strains expressing Achilles or GFP(S65T) under the control of a prestalk-specific *ecmAO* promoter (Williams *et al*., 1989). These strains also express either mScarlet-I (Bindels *et al*., 2016) or mRFPmars under *D19* promoter (Aubry and Firtel, 1999) whose activity is prespore-specific and is elevated earlier than the *ecmAO* promoter (Prabhu *et al*., 2007). mRFPmars is a monomeric RFP derived from *Discosoma* DsRed (Fischer *et al*., 2004) that is commonly used in *D. discoideum.* At 8 h of development, Achilles fluorescence appeared scattered in the aggregating stream (Fig. 4A) when GFP(S65T) fluorescence was barely detectable, except in a few cells (Fig. 4B). The fluorescence of mScarlet-I and mRFPmars was comparable at this streaming stage (Fig. 4C). Given that both Achilles and GFP(S65T) are expressed from an identical plasmid backbone, and that cells with green and red fluorescence appear mutually exclusive, the results suggest that Achilles allows for the earlier detection of cell-type bifurcation.

We further compared the expression of Achilles, GFP (S65T), mScarlet-I, and mRFPmars at later developmental stages under control of the *ecmAO* promoter. Three strains each carrying two fluorescent reporters were studied: Achilles/mRFPmars, GFP(S65T)/mRFPmars, and Achilles/mScarlet-I (Fig. 5). In early mound formation, before the appearance of the tip, Achilles and mScarlet-I as well as GFP(S65T) fluorescence became clearly detectable and appeared scattered throughout the mound (11 h, Fig. 5 A middle panel: 12 h, Fig. 5B middle panel: 12 h, Fig. 5C middle and lower panels). In contrast, the fluorescence of mRFPmars was not detected, except in a few cells (11 h, Fig. 5A, lower panel; 12 h, Fig. 5B, lower panel). By the time Achilles and mScarlet-I fluorescence appeared concentrated at the tip where prestalk cells accumulated, mRFPmars fluorescence became visible (12 h, Fig. 5A, lower panel; 13 h, Fig 5B, lower panel), suggesting that the level of mRFPmars fluorescence was similar to that of other FPs. From the tipped mound to the first finger stage (13–15 h), the fluorescence intensity of Achilles tendon in the anterior region was comparable to that of GFP(S65T) (13–14 h, Fig. 5 A middle panel: 14–15 h, Fig. 5B, middle panel). Similarly, the fluorescence intensities of mScarlet-I and mRFPmars were equally bright (13–14 h, Fig. 5 A lower panel: 14–15 h, Fig. 5B lower panel: 14–15 h, Fig. 5C lower panel). These results show that early detection of upregulated genes would benefit from the use of Achilles and mScarlet-I.

Although mScarlet, a brighter but slow-maturing sibling of mScarlet-I, has been used to label the cytosol and microtubules in *D. discoideum* (Paschke *et al*., 2018; Schweigel *et al*., 2022), we tested the suitability of mScarlet-I as a fusion tag and compared it with mCherry, an earlier-generation monomeric RFP variant with an emission peak at 610 nm (Shaner *et al*., 2004) also commonly used in *D. discoideum*. Under a confocal microscope using a 561 nm laser for excitation and bandpass filter at 595 nm, mScarlet-I as a standalone appeared uniform in the cytosol and nucleus of vegetative and slug cells (Fig. 6A). mScarlet-I fused to the Ras-binding domain of human Raf1 (RBD_hRaf1_), which preferentially binds to the GTP-bound form of Ras (Kae *et al*., 2004) shows clear localization in pinocytic cups (Fig. 6B). mCherry-RBD_hRaf1_ exhibited the same pattern, which is consistent with an earlier report (Veltman *et al*., 2016). mScarlet-I-RBD_hRaf1_ fluorescence was stronger than that of mCherry-RBD_hRaf1_ in vegetative cells, possibly because of the intrinsic brightness of mScarlet-I on top of the suboptimal excitation wavelength for mCherry (Fig. 6C). In the slug stage, in addition to the expected localization to the front cell–cell contact site (Fig. 6D) (Fujimori *et al*., 2019), a considerable number of aggregates were visible for mCherry-RBD_hRaf1_ but not for mScarlet-I-RBD_hRaf1_ (Fig. 6D). mScarlet-I-RBD_hRaf1_ fluorescence was approximately 4–5 times brighter than that of mCherry-I-RBD_hRaf1_ in both the anterior and posterior regions of the slug (Fig. 6E). Although mCherry has the advantage of having a longer wavelength that peaks at 610 nm, thus reducing the spectral overlap with yellow FP, the absence of aggregates makes mScarlet-I a top candidate red FP in *D. discoideum*.

### Expression of blue and large stokes shift FPs in *D. discoideum*

Next, we expand the color palette to include blue FPs. Two commonly used blue FPs are mTurquoise2: a blue FP derived from a Venus derivative Super Cyan Fluorescent Protein3A (SCFP3A) for high quantum yield and long fluorescence life-time (Goedhart *et al*., 2012) and TagBFP which is a monomeric blue FP derived from *Entacmaea quadricolor* TagRFP (Subach *et al*., 2008). In *D. discoideum*, only a few studies have used these FPs alone (Kundert *et al*., 2020; Paschke *et al*., 2019) or as a fusion tag (Mukai *et al*., 2016). Here, we tested further applicability of mTurquoise2 in addition to mTagBFP2 which is a brighter variant of TagBFP (Subach *et al*., 2011). As a standalone, mTurquoise2 and mTagBFP2 fluorescence appeared uniform in the cytosol and nucleus of vegetative cells (Fig. 7A, upper panels). Although this was also true in slug-stage cells for mTurquoise2 (Fig. 7A, left lower panels), aggregates were noticeable in mTagBFP2 (Fig. 6D: Fig. 7A, right lower panels). The results suggest that use of mTurquoise2 is preferable if aggregates are to be avoided during development.

When tagged to Lifeact, we observed strong mTurquoise2 fluorescence throughout the cell cortex of vegetative cells with marked concentration at the pinocytic cups and leading edges (Fig. 7B, left upper panel) in accordance with observations of F-actin binding mCherry-LimEΔcoil (Veltman *et al*., 2016). In contrast, mTagBFP2-Lifect fluorescence was localized to the rear of the cells (Fig. 7B; right upper panel). This difference was also observed in the slug-stage cells; mTurquoise2-Lifeact appeared uniformly in the cortex, whereas mTagBFP2-Lifeact was more concentrated in the cell rear (Fig. 7B, lower panels). As different actin-binding domains are known to show these localization differences (Lemieux *et al*., 2014), the results suggest that the fusion of these two FPs likely results in distinctly different affinities for F-actin. Other fusion tags were more forgiving regarding the choice of FPs. The PH domain of Akt/PKB (PH_Akt_) fused to mTurquoise2 or mTagBFP2 showed a localization pattern similar to that of PH_Akt_-Achilles for both vegetative and slug-stage cells, with mTurquoise2 exhibiting a higher brightness (Fig. 7C; Fig. 2A, C, and D). They appeared mostly in the pinocytic cups of vegetative cells and at the cell front of the slug. HistoneH1-mTurquoise2 and HistonH1-mTagBFP2 also showed proper localization to the nucleus in vegetative and slug cells (Fig. 7D). These data demonstrate that mTurquoise2 is well expressed as a standalone fusion protein in the single- and multicell stages of *D. discoideum.* mTagBFP2 may be less bright and its localization may be more sensitive to the protein tag.

Next, we explored the use of LSSmGFP, a large stokes shift variant of the hyperfolder YFP engineered using a top-down structure-based approach for 405 nm excitation (Campbell *et al*., 2022). Since green fluorescence emission under 405 nm excitation is not commonly utilized in *D. discoideum*, we first evaluated autofluorescence in the wild-type Ax4 background. We found no significant autofluorescence except for some small particles whose mean intensity was approximately 5% above the background (Fig. 8A and B). When vegetative cells expressing LSSmGFP under *act15* promoter were excited at 405 nm, green fluorescence appeared uniformly in the cytosol, with an average intensity exceeding 150% above the background (Fig. 8C and D). Spectral bleed-through of LSSmGFP fluorescence at 447 nm was undetectable, as determined by comparison with Ax4. When excited at 488 nm, there was minor fluorescence above the background, the intensity of which was approximately 1% of that observed under 405 nm excitation (Fig. 8C and D 488 Ex). The level of cross-excitation was consistent with the reported spectral properties of LSSmGFP (Campbell *et al*., 2022).

Next, we tested the applicability of LSSmGFP to live-cell imaging in *D. discoideum*. Standalone LSSmGFP showed uniform fluorescence in both vegetative and slug-stage cells (Fig. 8E). LSSmGFP-Lifeact was localized to the cell cortex, pinocytic cups in vegetative cells, and leading edges in slug-stage cells (Fig. 8F), similar to mTurquoise2-Lifeact (Fig. 7A). Fluorescence of LSSmGFP-HistoneH1 was restricted to the nucleus, as expected, in both vegetative and slug-stage cells (Fig. 8G). Next, we tested 3-color imaging by constructing a strain that co-expressed GcvH1-mTagBFP2, LSSmGFP-HistoneH1 and PKBR1(N150)-Achilles (Fig. 8H). GcvH1(N99) is the N-terminal mitochondrial localization sequence of GcvH1 gene (Perry et al 2020). PKBR1(N150) is the N-terminal sequence of PKBR1 that includes a myristoylation signal and labels the plasma membrane (Meili *et al*., 2000). While these probes showed the expected localization (Fig. 8H; vegetative cells), we noticed with 405 nm excitation some bleed-through from the fluorescence of GcvH1(N99)-mTagBFP2 to the 525 nm emission range (Fig. 8H, cyan and green). While there is some overlap with autofluorescence, the results demonstrate that LSSmGFP is a viable option for multi-color imaging in *D. discoideum*.

### Near-infrared fluorescence imaging in *D. discoideum*

For NIR fluorescence imaging, iRFP and mIFP derived from *Rhodopseudomonas palustris* and *Bradyrhizobium sp.,* respectively have been employed in *D. discoideum* as a standalone (Kundert *et al*., 2020; Ohta *et al*., 2018). However, these have limited applications owing to their low brightness, dimerization, and relatively large size. Monomeric NIR FP derived from bacterial phytochrome photoreceptors (Shcherbakova *et al*., 2015) may overcome these drawbacks. These FPs consist of <u>P</u>er-<u>A</u>RNT-<u>S</u>im (PAS) and c<u>G</u>MP-specific phosphodiesterase, <u>a</u>denylyl cyclase, and <u>F</u>hla (GAF) domains, which contain the chromophore biliverdin (Rumyantsev *et al*., 2015; Wagner *et al*., 2007). Notably, the truncated variant, miRFP670nano, consisted of only a single GAF domain. Its size (∼17 kDa) is close to half that of GFP (27 kDa) and has high molecular brightness (Oliinyk *et al*., 2019), making it a prime candidate for use in *D. discoideum.* Here, we prepared cells expressing miRFP670nano3 (Oliinyk *et al*., 2022)—a brighter variant of miRFP670nano, under *act15* promoter. Cells were incubated in culture medium with biliverdin and developed on an agar plate containing 50 µg/mL biliverdin (Kundert *et al*., 2020). We observed that in vegetative cells as well as in cells within slugs, miRFP670nano3 fluorescence appeared uniform in the cytosol and nucleus (Fig. 9A). HistoneH1-miRFP670nano3 was localized in the nucleus (Fig. 9B). PH_Akt_-miRFP670nano3 was localized to the pinocytic cup in vegetative cells (Fig. 9C, upper panel) and cell–cell contact sites in slug-stage cells (Fig. 9C, lower panel). As expected, when miRFP670nano3 was expressed under the prespore-specific promoter, miRFP670nano3 signals were detected in the posterior region of the slugs (Fig. 9D). When imaging a fruiting body, caution must be taken to account for weak autofluorescence in the prestalk and stalk regions, whose intensity is approximately 35% of the background (Fig. S3A–C). This may not be an issue when the signal is an order of magnitude higher than the background fluorescence, as in the case of miRFP670nano3 expressed under the control of the D19 promoter (Fig. S3D–F).

Multi-color imaging may also be deployed using the self-labeling protein HaloTag, a modified haloalkane dehalogenase that covalently binds to a chloroalkane linker substrate (Los *et al*., 2008). In *D. discoideum*, HaloTag-fused PTEN and GFP-tagged PH domains of Akt/PKB have been used together with the red fluorescent TMR-conjugated Halo-ligand to track protein and lipid dynamics in the plasma membrane (Arai *et al*., 2010; Matsuoka *et al*., 2012). Since the spectral properties depend on the fluorophore that labels the cell-permeable ligand, one may take advantage of choosing the type of fluorophore and its dosage to avoid spectral overlap with FPs and overstaining (Grimm *et al*., 2017). Here, we tested the silicone rhodamine-based label SaraFluor 650T, which was excited at 650 nm to obtain NIR fluorescence (Yanagawa *et al*., 2018). After labeling (see Supplementary information), vegetative cells expressing HaloTags showed uniform signals in the cytosol (Fig. 9E, upper panel). Although the cells were labeled only during growth, fluorescence was still visible in the slugs (Fig. 9E, lower panel). PH_Akt_-Halo and HistoneH1-Halo showed proper localization patterns in both vegetative and slug-stage cells (Fig. 9F and G). These results demonstrate that HaloTag in combination with the ligand SaraFluor 650T provides an alternative to NIR FPs in *D. discoideum*.

### Four-color live-cell fluorescence imaging in *D. discoideum*

We envisaged that the use of blue and NIR fluorescence in combination with the green and red FP tested above, owing to their brightness and relatively good spectral separation, would render multi-color imaging practical in *D. discoideum*. We first imaged four FPs, namely, GcvH1(N99)-mTagBFP2, Dajumin-mScarlet-I, PKBR1(N150)-Achilles, and HistoneH1-miRFP670nano3, which target the mitochondria (Perry *et al*., 2020), plasma membrane (Meili *et al*., 2000), contractile vacuole (Gabriel *et al*., 1999) and nucleus (Nagasaki *et al*., 2002), respectively. We used three plasmids with three selection markers, one of which carried two FP-encoding genes (Supplementary Information, Table S2) to circumvent the limited number of selection drugs. In vegetative cells, we observed the expected localization patterns in organelles and plasma membranes, with good separation between the respective channels (Fig. 10A, Movie S1). Live-cell imaging revealed the dynamics and positional relationships between these structures. Mitochondria were dispersed in regions free of contractile vacuoles and nuclei (Fig. 10A). Contractile vacuoles showed sequential deformation (Fig. 10A, white arrows). The nucleus continued to change its position between an area adjacent to the plasma membrane (Fig. 10A, 0–50 s) and a more distal area (Fig. 10A, white arrow, 60–70 s), appearing as though it was being pushed away by a contractile vacuole. We also noticed that Dajumin-mScarlet-I fluorescence appeared weakly in the plasma membrane (Fig. 10A), which is in contrast to an earlier report of Dajumin-GFP showing stricter localization to the contractile network (Gabriel *et al*., 1999).

Next, we imaged five FPs: LSSmGFP-Lifeact, GcvH1(N99)-mTagBFP2, Golvesin-Achilles, PKBR1(N150)-mScarlet-I and HistoneH1-miRFP670nano3. These target the F-actin, Golgi apparatus (Schneider *et al*., 2000), plasma membrane, and nucleus. To obtain a strain that expressed all five FPs, the PKBR1(N150)-mScarlet-I-2x sequence was first inserted into the *act5* locus (Paschke *et al*., 2018) by homologous recombination, followed by the introduction of other FPs into extrachromosomal vectors (see Supplementary information). Fig. 10B shows representative time-lapse images of the cells dissociated from the slugs. Here, LSSmGFP-Lifeact and GcvH1(N99)-mTagBFP2 images were first obtained by 405 nm excitation followed by sequential acquisition of fluorescence images in the yellow, red, and NIR spectra by excitation at 488, 560, and 640 nm, respectively. To isolate LSSmGFP fluorescence in the green channel, weak cyan fluorescence originating from 405 nm-excited GcvH1(N99)-mTagBFP2 was estimated from other channels and subtracted. To isolate Achilles fluorescence in the yellow channel, weak cyan and green fluorescence from 405 nm-excited GcvH1(N99)-mTagBFP2 and 488 nm-excited LSSmGFP-Lifeact were estimated and subtracted (see Supplementary information). Although more thorough spectral unmixing should be employed for rigorous signal separation (Leiwe *et al*., 2024), subtracting these two components results in well-isolated LSSmGFP and Achilles signals. While there was noticeable cell–cell heterogeneity in the fluorescence intensity, for cells that appeared positive in all 5-channels, the FP tags exhibited the expected intracellular localization in the respective emission spectra. Golvesin-Achilles appeared not only in the perinuclear region, as reported for Golvesin-GFP (Schneider *et al*., 2000) but also in the ER, likely because of the brightness and fast maturation of Achilles. This allowed observation of the movement of the nucleus and mitochondria, which are tightly embedded in the adjacent ER network (Fig. 10B and Movie S2). Four-color time-lapse imaging was performed using PH_Akt_-mTurquoise2 (PIP3), mCherry-RBD_hRaf1_ (Ras-GTP), CRIB_PakB_-Achilles (Rac-GTP), and Halo-Lifeact (F-actin) (Supplementary Text, Fig. S4, Movies S3 and S4), demonstrating the applicability of multi-color imaging to *D. discoideum*. While the localization patterns of FP-tagged proteins must be further scrutinized by other means, such as immunostaining, these results demonstrate the relative feasibility of multi-color fluorescence imaging for the detailed analysis of collective cell migration in *D. discoideum*.

### Combining cellulose-staining and FPs in *D. discoideum*

Fluorescent dyes may also be useful in multichannel imaging. In fixed *Dictyostelium* samples, calcofluor white is conventionally used to label cellulose, which is a major component of the extracellular matrix, referred to as the slime sheath, stalk tube, surface of the mature stalk, and spore cells in the fruiting bodies (Blanton *et al*., 2000; Yamada and Schaap, 2019). By including calcofluor in the agar substrate, thereby allowing it to diffuse and be absorbed in the sample, we found that it could be utilized for live-cell imaging (Fig. 11A, Movie S5). Furthermore, we observed that Direct Fast Scarlet 4BS (Pontamine Fast Scarlet 4 B), a carbohydrate-binding dye used in plants to visualize cellulose at an excitation wavelength of 561 nm (Anderson *et al*., 2010) also stained the stalk region of slugs and the fruiting body without affecting development (Fig. 11B). The staining patterns of the two dyes are indistinguishable. In the example shown in Fig. 11C, the stalk was stained with calcofluor white, and two prestalk subtypes, pstAO and pstB, and prespore cells were visualized using FPs in the fruiting body (Fig. 11C).

## Summary and Discussion

Here, we demonstrated that Achilles and mScarlet-I are two-color alternatives for live-cell imaging of *D. discoideum* both in the vegetative and development stages. Along with brightness, their rapid maturation facilitates the early detection of prestalk-specific gene expression. These characteristics, along with the degradation tag, aids in addressing temporal and spatial patterns of developmental gene expression. This study expands the FP color palette to include the blue and NIR spectra. For blue FP, mTurquoise2 was the protein of choice for *D. discoideum* because of its high fluorescence and absence of aggregates. While blue FPs have not been well adopted, our results indicate that mTurquoise2 can be considered for practical use, providing access to microscopes with high-quantum-efficiency photomultipliers or sCMOS cameras. Although mTagBFP2 showed some aggregation and interference in the localization of tagged proteins (Lee *et al*., 2013; Vještica *et al*., 2020), its use is worth considering when the emission spectrum is more compatible with the DAPI filter set. Regarding NIR imaging, our data indicated that miRFP670nano3 was sufficiently bright, did not aggregate, and exhibited minimal interference with protein localization in *D. discoideum*. It should be noted that, under our optical setup, the duration of excitation and signal detection had to be extended by approximately five-fold or more compared with those required for other FPs to achieve a comparable S/N ratio. Owing to this limitation, NIR FP should be employed for tagging strongly expressed proteins, reserving brighter FPs for those with weak expression. In the future, this limitation may be partially overcome by introducing the phycocyanobilin synthesis system (SynPCB) (Sakai *et al*., 2021) as an alternative to exogenous biliverdin.

Using these two emission spectra, we demonstrated the feasibility of four-color fluorescence imaging using confocal microscopy. Using a conventional confocal microscope, image acquisition can be performed serially in tandem or accelerated, for example, by repeating 2-laser excitation 2-emission readout serially, provided that appropriate dichroic mirrors and filter sets are available. Given the bleed-through of LSSmGFP to 525 nm with 405 nm excitation, a combination of FPs for practical application in multi-color confocal microscope imaging without requiring spectral unmixing is mTurquoise2, Achilles, mScarlet-I, and HaloTag with the NIR ligand. Further assessment of spectral unmixing is necessary if five or more emission spectra are required.

Another challenge for deploying multi-color fluorescence imaging in *D. discoideum* is the choice of vehicles on which genetically encoded FPs and tags are introduced. Here, extrachromosomal vectors that use the origin of replication of naturally occurring plasmids (Veltman, Akar, *et al*., 2009) were employed. Provided that the tags are small enough, one may choose to have a single plasmid carrying two fusion tags (Supplementary Information), as we did here with GcvH1(N99)-mTagBFP2 and HistoneH1-miRFP670nano3 (Fig. 10), and PH_Akt_-mTurquoise2 and Halo-Lifeact (Fig. S4). This allowed the expression of four tags with three commonly available selection markers in *D. discoideum*: G418, Blasticidin and Hygromycin. However, cells expressing all four probes equally at high levels were rare, likely because of heterogeneity in the plasmid copy number. Furthermore, maintenance of these plasmids requires the constant selection of multiple drugs, potentially affecting cell physiology. These caveats may be mitigated by knock-in strategies so that all tags are single-copy and the resulting cell lines may be stable in the absence of the selection drug (Paschke *et al*., 2018). *rps30* locus in the duplicated region of chromosome 2 of the Ax3 and Ax4 strains (Eichinger *et al*., 2005) and *act5* locus, which is redundant among 17 actin genes (Joseph *et al*., 2008) have been shown to be useful as a “safe harbor” (Corrigan and Chubb, 2014; Muramoto *et al*., 2010). Here, we employed a knock-in at *act5* locus to introduce an FP expression cassette and three extrachromosomal vectors to express four FPs for 5-color imaging. Exploring additional safe harbors should facilitate applications where the homogeneous expression of multiple FPs is important. The disadvantage of this approach is the reduced brightness due to the lower expression level and relatively laborious process of generating a knock-in. Other methods include the use of randomly integrated plasmids (Fey *et al*., 1995) which should also be considered for a high copy number.

## Acknowledgments

We thank Douwe Veltman for pDM304, pDM323, pDM324, pDM326, pDM340, pDM358 (National BioResource Project Nenkin plasmid; G90008, G90073, G90074, G90067, G90063, G90009), pDM1208, pDM1209 and pDM1489, Chris Thompson for p*ecmO*:labile-GFP, Jonathan Chubb for providing mNeonGreen vector, Robert Kay for pDM1501 (Addgene plasmid; #108997), Atsushi Miyawaki and the RIKEN BRC for providing Achilles through the National BioResource Project of the MEXT, Japan. The plasmids constructed in this study are available from the NBRP-Nenkin stock center (https://nenkin.nbrp.jp/).

## Abbreviations

FP: fluorescent protein
GFP: green fluorescent protein
RFP: red fluorescent protein
CFP: cyan fluorescent protein
YFP: yellow fluorescent protein
NIR: near-infrared
PIP3: phosphatidylinositol (3,4,5) trisphosphate
PH: Pleckstrin homology
RBD: Ras-binding domain
CRIB: Cdc42/Rac interactive binding.

