## Supplementary Texts and Figures for "Multi-color fluorescence live-cell imaging in *Dictyostelium discoideum*"

### Supplementary information for Multi-color fluorescence live-cell imaging in *Dictyostelium discoideum*

#### Supplementary methods

#### Plasmid construction

The plasmids and primers used are listed in Table S2. The plasmids constructed in this study will be available through NBRP-nenkin (https://nenkin.nbrp.jp/). To obtain N- and C-terminus FP tag vectors, Dicty-codon optimized DNA for mTagBFP2 (Subach et al. 2011), mTurquoise2 (Goedhart *et al.*, 2012), Achilles (Yoshioka-Kobayashi *et al.*, 2020), mScarlet-I (Bindels *et al.*, 2016), miRFP670nano3 (Oliinyk *et al.*, 2022) and LSSmGFP (Campbell *et al.*, 2022) were synthesized and inserted into Bgl II or Spe I sites of pDM304, pDM326 or pDM358 (Veltman, Akar, *et al.*, 2009) by fusing PCR generated inserts and linearized vectors with In-Fusion enzyme (In-Fusion Snap Assembly cloning kit, TAKARA) or by ligation with T4 DNA ligase (Ligation high Ver. 2, TOYOBO). PCR amplified fragments were cloned into pCR Blunt II-TOPO vector (Zero Blunt TOPO PCR Cloning Kit, Invitrogen) before digestion for ligation. To obtain N- and C-terminus HaloTag vectors, the coding sequence of HaloTag was PCR amplified from pFC14A_HaloTag CMV Flexi Vector (Promega) and inserted into Bgl II or Spe I sites of pDM304 with In-Fusion enzyme or T4 DNA ligase. For tagged FPs, coding sequences of HistoneH1, PKBR1 (1–150 a.a containing myristoylation motif (Meili *et al.*, 2000)), Dajumin (Gabriel *et al.*, 1999), Akt/PKB (1– 111 a.a, containing PH domain (Meili *et al.*, 1999)), PakB (338– 431 a.a, containing CRIB domain (Veltman *et al.*, 2012)), GcvH1 (1-33 a.a containing mitochondrial localization signal (Perry *et al.*, 2020)) and Golvesin (Schneider *et al.*, 2000) were PCR amplified using Phusion polymerase (New England Biolab) from genomic DNA or cDNA. A hygromycin resistance cassette was digested from pDM1489 (Paschke *et al.*, 2018) and inserted into the XhoI and BamHI sites of pDM1208 to obtain a hygromycin-resistant expression vector #HH67. The Ras-binding domain of human Raf1 were digested from pDM358_mRFPmars-Raf1RBD (Nakajima *et al.*, 2014) by BglII and SpeI. These fragments were cloned and inserted into the BglII and SpeI cloning sites of expression vectors. Similarly, annealed oligos of Lifeact (Riedl *et al.*, 2008) were inserted into the expression vectors at the BglII and SpeI cloning sites.

The *V18*, *ecmAO*, *ecmB* and *D19* promoters (Ceccarelli *et al.*, 1991; Early *et al.*, 1993; Early and Williams, 1989; Singleton *et al.*, 1989) were cloned from genomic DNA. The promoter region of *coaA* was PCR-amplified from pDM1209 (Paschke *et al.*, 2018). The cloned fragments were inserted into the FP expression vectors described above at the BglII and XhoI sites. To generate an mRFPmars-expressing vector with the *ecmAO* and *D19* promoter, the promoter element and coding sequence of mRFPmars were amplified from pEcmAO-mRFPmars and pD19-RFP (Fujimori *et al.*, 2019), respectively, and the cloned fragments, including the promoter and coding sequences, were inserted into the XhoI and SpeI sites of pDM358.

To obtain dual FP expression vectors #HH143 for Halo-Lifeact and $PH_{Akt}$-mTurquoise2 and #HH504 for HistoneH1-miRFP670nano3 and GcvH1(N99)-mTagBFP2 (Table S2), the fragments containing the promoter region, coding sequence of the fluorescence reporter and the terminator region, were PCR amplified from #HH114 and #HH499 and inserted into the NgoMIV site of #HH76 and #HH500, respectively. To generate labile-Achilles, the N-terminal Ubi sequence excised from p*ecmO*:labile-GFP by BglII and BamHI was fused to the N-terminus of Achilles at the BglII site of pDM304_*V18*p:Achilles (pDM304_*V18*p:Labile-Achilles). GFP cassette on pDM340 (Veltman, Keizer-Gunnink, *et al.*, 2009) was replaced by Achilles to generate a Dox-inducible Achilles expression vector pDM340-Achilles. To generate pDM1501_PKBR1(N150)-mScarlet-I-2x, the PKBR1(N150)-mScarlet-I fragment was amplified and inserted into the BglII and SpeI sites of pDM1501 (Paschke *et al.*, 2018), and an additional mScarlet-I fragment was inserted into the SpeI site located between PKBR1(N150) and mScarlet-I.

#### Generation of the 4- and 5-color fluorescence-labeled cell line

For 4-color imaging (Fig. 10A and Fig. S4), single vectors (G418-resistance) carrying two tags, namely HistoneH1-miRFP670nano3 and GcvH1(N99)-mTagBFP2 (#HH504), or Halo-Lifeact and $PH_{Akt}$-mTurquoise2 (#HH143), were employed together with the Hygromycin- and Blasticidin-resistant vectors. For 5-color imaging (Fig. 10B), we generated a knock-in of mScarlet-I-tagged PKBR1(N150) at the *act5* locus and transformed it with three plasmids, one of which harbored two FP-tagged genes. Following a previous study (Paschke *et al.*, 2018), we used the *act5 locus* to insert PKBR1(N150)-mScarlet-I-2x. PKBR1(N150) was fused to two tandem repeats of mScarlet-I to increase brightness. The sequence encoding the tagged protein with flanking loxP sites was inserted immediately after the *act5* promoter, together with a hygromycin resistance cassette (Paschke *et al.*, 2018). Cells were electroporated with 10 μg of 5.5 kbp fragment of #HH571 pDM1501-PKBR1(N150)-mScarlet-I-2x by excising with NgoMIV. Cells were selected in the presence of 60 μg/ml Hygromycin B as described earlier (Paschke *et al.*, 2018), and clones with less cell-cell heterogeneity in the fluorescence signals were chosen. The hygromycin resistance cassette was then removed from the locus by transiently expressing Cre-loxP (Faix *et al.*, 2004) for the expression of another FP using an extrachromosomal vector with a hygromycin cassette. The strain was transformed with three plasmids, one of which harbored two FP-tagged genes. The expression plasmids for LSSmGFP-Lifeact (#HH605), HistoneH1-

miRFP670nano3/GcvH1(N99)-mTagBFP2 (#HH501), and Golvesin-Achilles (#HH497) were used (Table S2).

#### Cell labeling for NIR imaging and cellulose imaging

For miRFP670nano3 expressing cells, cells were pre-incubated overnight in growth medium with biliverdin (Sigma-Aldrich) at the final concentration of 10 or 50 µg/ml before harvesting. For observation of slug and fruiting body, cells were developed on the agar plate contained 50µg/ml biliverdin. To obtain HaloTag-labeled cells, vegetative cells expressing HaloTag-fusion protein were washed once and suspended in 400 uL PB with 1 µM (final conc.) Sarafluor 650T ligand (Goryo Chemical) at cell density of $10^7$ cells/ml and shaken at 22 for 30 minutes. Labeled cells were washed thrice and resuspended in PB (0.5 mL) at the same cell density. The cells were then incubated for 30 min in a shaken tube and washed twice with PB.

For cellulose staining, the agar plate contained 0.1 or 1 mg/ml of Fluorescent Brightener 28 (Calcofluor white, MP Biomedicals 158067) or 1 mg/ml of Direct Fast Scarlet 4BS (FUJIFILM WAKO 043-28272) (Anderson *et al.*, 2010). For Direct Fast Scarlet 4BS, agar was dissolved in Bonner's Salt Solution (0.6 g NaCl, 0.75 g KCl, 0.3 g $CaCl_2$ in 1 L water: (Bonner and Savage, 1947)). Stock solution of Fluorescent Brightener 28 and Direct Fast Scarlet 4BS were dissolved at the concentration of 10 mg/ml in MilliQ water and PBS, respectively and stored in dark at 4 °C.

#### Calibration of breed-through for 5-color fluorescence imaging

For the 5-FP expressing strain (Fig. 10B), LSSmGFP fluorescence was separated from the spectral bleed-through of mTagBFP2, and Achilles fluorescence was corrected for the cross-excitation of LSSmGFP. To this end, the images were first background-corrected based on the fluorescence of the parental Ax4 strain (Fig. 8A). Binarized masks of autofluorescent vesicles were generated by manually thresholding the Ex 488 nm and Em 525/50 nm images of the Ax4 cells. These regions were removed from the Ax4 and LSSmGFP/Ax4 cell masks before computing the mean background fluorescence intensities in each channel (Fig. 8 B and D; both n = 6 cells). The background was subtracted from each channel to obtain images of the 5-FP expressing cells.

The background-subtracted images were further processed to remove cross excitons and bleed-through. Based on the LSSmGFP/Ax4 images, the ratio ($\alpha$) between the Ex 405 nm, Em 525/50 nm images and the Ex 488 nm, Em 525/50 nm images was obtained. To obtain the corrected Golvesin-Achilles images in the 5-FP expressing cells (Fig. 10B), Ex 405 nm, Em 525/50 images ($I_{405ex,\ 525em}$) were multiplied by $\alpha$ and subtracted from the Ex 488 nm, Em 525/50 nm images ($I_{488ex,\ 525em}$). Next, the bleed-through from the 405 nm excited GcvH1(N99)-mTagBFP2 fluorescence into the green (525/50 nm) channel was removed. Six 110 × 90 μm areas were selected from an Ex 405 nm, Em 447/60 nm image of a 5-FP expressing cell to obtain masks of the mitochondrial marker GcvH1(N99)-mTagBFP2 by binarization using IsoData algorithm. The average fluorescence intensity ratio ($\beta$) of the mitochondrial masks at Ex 405, Em 447 nm and Ex 405 nm, Em 525/50 nm was computed (6 areas). To obtain LSSmGFP-Lifeact images, Ex 405 nm, Em 447/60 nm images ($I_{405ex,\ 447em}$) were multiplied by $\beta$ and subtracted from the Ex 405 nm, Em 525/50 nm images ($I_{405ex,\ 525em}$).

#### Supplementary text

#### 4-color imaging of PIP3, small GTPases and F-actin.

For 4-color imaging of F-actin and its regulators, we constructed a strain that expressed markers for Ras-GTP, PIP3, Rac-GTP, and F-actin. Activated Ras was labeled with mCherry-$RBD_{hRaf1}$. PIP3 was detected using $PH_{Akt}$-mTurquoise2. The localization of activated Rac was visualized using Achilles fusion to the CRIB domain of PakB ($CRIB_{PakB}$), which preferentially binds to the Rac-GTP form (Veltman *et al.*, 2012). F-actin was visualized by labeling the expressed Halo-Lifeact using the HaloTag ligand, SaraFluor 650. Figure S4A and Movie S3 show representative time-lapse images of vegetative cells expressing the above four FP. These localization patterns agreed well with an earlier observations based on mCherry-LimE Δ coil (Veltman *et al.*, 2016). As expected, Halo-Lifeact appeared in the cell cortex at a marked concentration in the pinocytic cup, similar to mTurquoise2-Lifeact (Fig. 7B: Fig. S4A). $PH_{Akt}$-mTurquoise2, $CRIB_{PakB}$-Achilles, and mCherry-$RBD_{hRaf1}$ showed more selective localization in the pinocytic cup than in Halo-Lifeact (Fig. S4A), consistent with earlier observations based on GFP- and mCherry-fused probes (Rupper *et al.*, 2001; Veltman *et al.*, 2016). Subtle differences were observed between $PH_{Akt,}$ $CRIB_{PakB}$ , $RBD_{hRaf1}$ and Lifeact. $CRIB_{PakB}$-Achilles was weakly localized to the F-actin cortex, marked with Halo-Lifeact; however, this was not as obvious for $PH_{Akt}$-mTurquoise2 and mCherry-$RBD_{hRaf1}$ (Fig. S4A). In addition, there was a small protrusion where $CRIB_{PakB}$-Achilles and Halo-Lifeact showed strong localization, whereas $PH_{Akt}$-mTurquoise2 and mCherry-$RBD_{hRaf1}$ did not (Fig. S4A; 0 s, white arrows). The slight difference in the observed patterns of Ras-GTP and Rac-GTP in vegetative cells is in line with earlier observation of GFP-$RBD_{hRaf1}$ (GFP-RBD) and $CRIB_{PakB}$-mCherry (RFP-PakB-CRIB): Rac-GTP at the rim of the pinocytic cup and its edge, and Ras-GTP only at the cup rim (Buckley *et al.*, 2020). $PH_{Akt}$-mTurquoise2 also appeared in an internalized macropinosome (Fig. S4A, 0–30 s), whereas $CRIB_{PakB}$-Achilles and mCherry-$RBD_{hRaf1}$ did not.

Furthermore, we checked the intracellular localization of FP-tagged proteins in slug stage cells (Fig. S4B and Movie S4). Although there was large cell-cell heterogeneity at the level of FP expression, we were always able to find a few cells in a field of view that expressed all four FPs. In the representative example shown, which highlights the migrating prespore cells in a slug, F-actin appeared densely at several protrusions, mainly located on the front side of the cell (Fig. S4B, Halo-Lifeact). The PIP3 marker $PH_{Akt}$-mTurquoise2 appeared at the cell-cell contact site, in accordance with earlier reports based on $PH_{Akt}$-GFP and $PH_{CRAC}$-GFP (Dormann *et al.*, 2002; Fujimori *et al.*, 2019; Hashimura *et al.*, 2019). Similar localization patterns were observed for the Ras-GTP marker mCherry-$RBD_{hRaf1}$. $CRIB_{PakB}$-Achilles on the other hand appeared to be broadly distributed throughout the cell front (Fig. S4B; 10 s, white arrows), hinting at separate roles that Ras/PIP3 and Rac play in promoting the cell-cell contact (Fujimori *et al.*, 2019) and forward migration.

#### Supplementary Figures

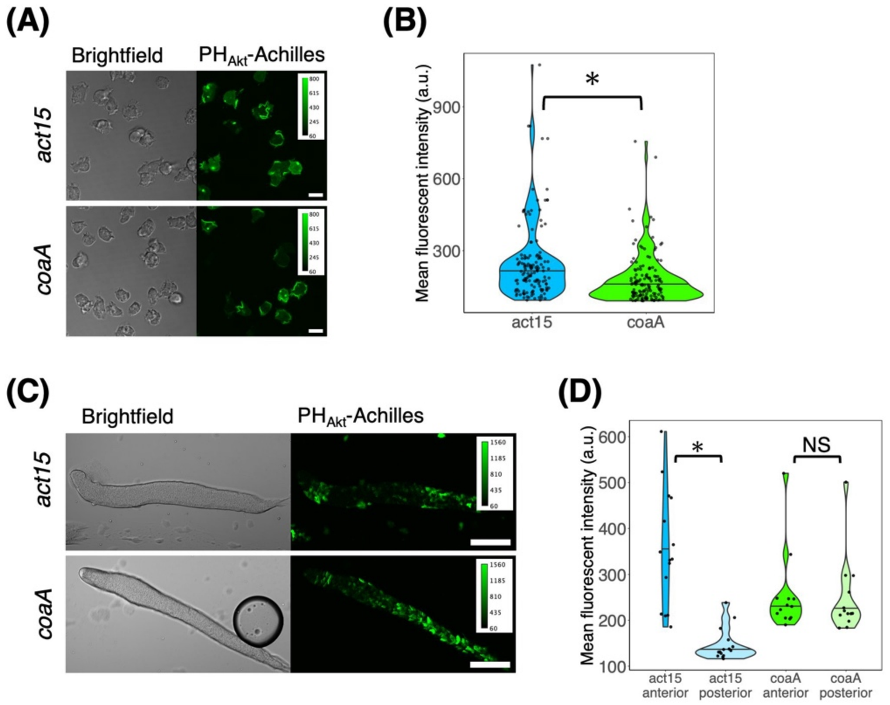


**Fig. S1. Comparison of $PH_{Akt}$-Achilles expression under *coaA and act15* promoter.**

(A) Cells carrying *act15p*:$PH_{Akt}$-Achilles (upper panels) and *coaAp*:$PH_{Akt}$-Achilles (lower panels) (left: transmitted light, right: green channel). Vegetative-stage. Scale bar, 10 µm. (B) Violin plot of the single-cell mean fluorescent intensity of cells carrying *act15p*:$PH_{Akt}$-Achilles and *coaAp*:$PH_{Akt}$-Achilles. Vegetative-stage (*act15*p:$PH_{Akt}$-Achilles, n = 91 cells. *coaAp*:$PH_{Akt}$-Achilles, n = 143 cells). The black line indicates the median. *: $P < 10^{-3}$. (C) Representative snapshots of slugs (*act15p*:$PH_{Akt}$-Achilles, upper panel. *coaAp*:$PH_{Akt}$-Achilles, lower panels) . Scale bar, 100 µm. The anterior-posterior axis of the slug is from left to right. (D) Violin plot of the mean fluorescent intensities of the anterior and the posterior region (*act15p*:$PH_{Akt}$-Achilles, n = 15 slugs. *coaAp*:$PH_{Akt}$-Achilles, n = 13 slugs). The black line indicates the median value.*: $P < 10^{-6}$. NS: not significant ($P > 0.05$).

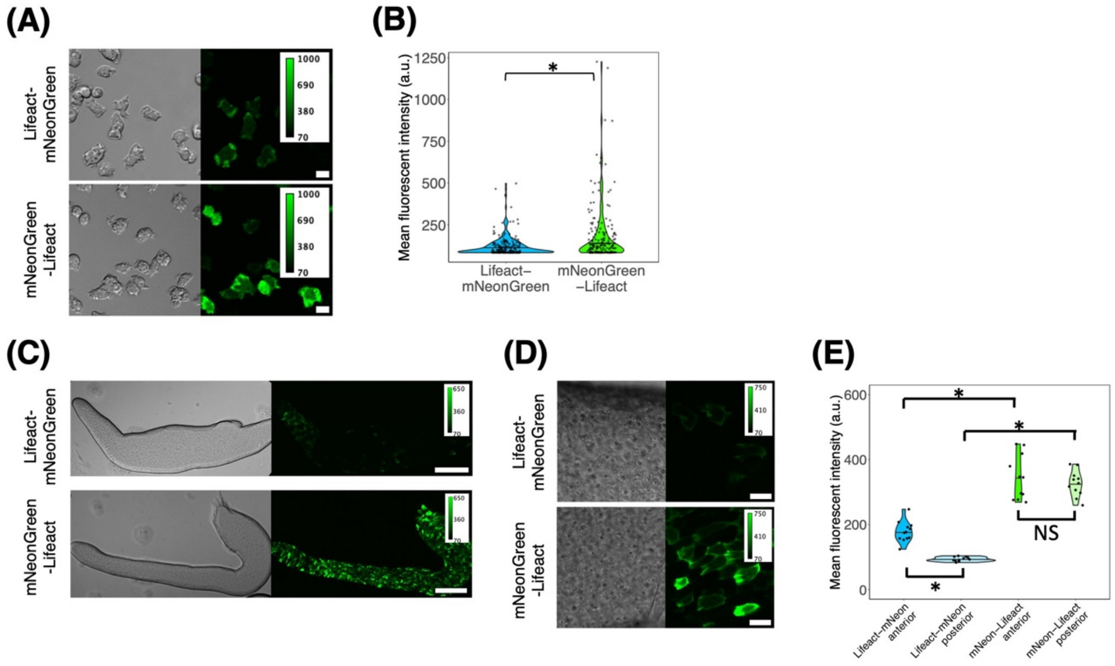


**Fig. S2. Fluorescence of Lifeact-mNeonGreen was decreased at the slug stage in *D. discoideum* cells.**

(A) Cells carrying *act15p*:Lifeact-mNeonGreen (upper panels) and *act15p*:mNeonGreen-Lifeact (lower panels) (left: transmitted light, right: green channel). Scale bar, 10 µm. (B) Violin plot of the single-cell mean fluorescent intensity of Lifeact-mNeonGreen and mNeonGreen-Lifeact expressing cells. Vegetative-stage (*act15p*:Lifeact-mNeonGreen, n = 191 cells. *act15p*:mNeonGreen-Lifeact, n = 191 cells). The black line indicates the median. *: $P < 10^{-4}$. (C) Representative snapshots of slugs (*act15p*:Lifeact-mNeonGreen, upper panels; *act15p*:mNeonGreen-Lifeact, lower panels). Scale bar, 100 µm. (D) High magnification images of slug expressing Lifeact-mNeonGreen (upper panels) and mNeonGreen-Lifeact (lower panels). Scale bar, 10 µm. The anterior-posterior axis of the slug is from left to right. (E) Violin plot of Lifeact-mNeonGreen and mNeonGreen-Lifeact fluorescent intensities of the anterior and the posterior region (Lifeact-mNeonGreen, n = 12 slugs. Lifeact-mNeonGreen or mNeonGreen-Lifeact, n = 11 slugs). The black line indicates the median. *: $P < 10^{-5}$. NS: not significant ($P > 0.05$).

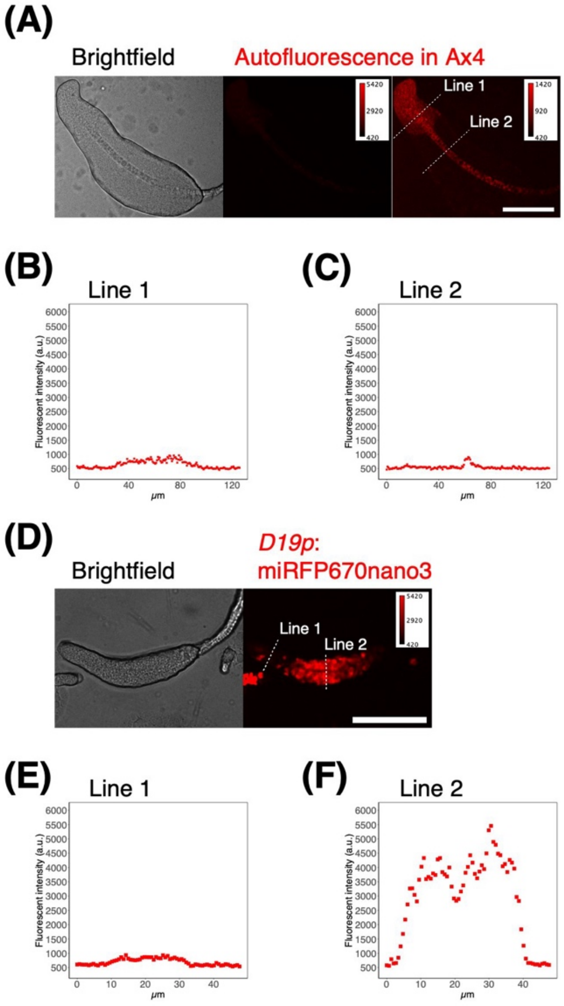


**Fig. S3. Autofluorescence appears in prestalk and stalk region of the fruiting body with 638 nm excitation in *D. discoideum* cells.**

(A) The autofluorescence of Ax4 cells during the culmination stage (638nm excitation and 685/40 emission). The brightfield image (left panel), autofluorescence (middle) and autofluorescence after contrast adjustment (right). Scale bar, 100 µm. (B, C) Line profile of the autofluorescence intensity along with the line 1 (B) and line 2 (C) in (A). (D) Snapshots of a culminant (*D19p*:miRFP670nano3; 638nm excitation and 685/40 emission). The brightfield (left) and fluorescence image (right). Scale bar, 100 µm. (E, F) Line profile of the autofluorescence intensity along with the line 1 (E) and line 2 (F) in (D).

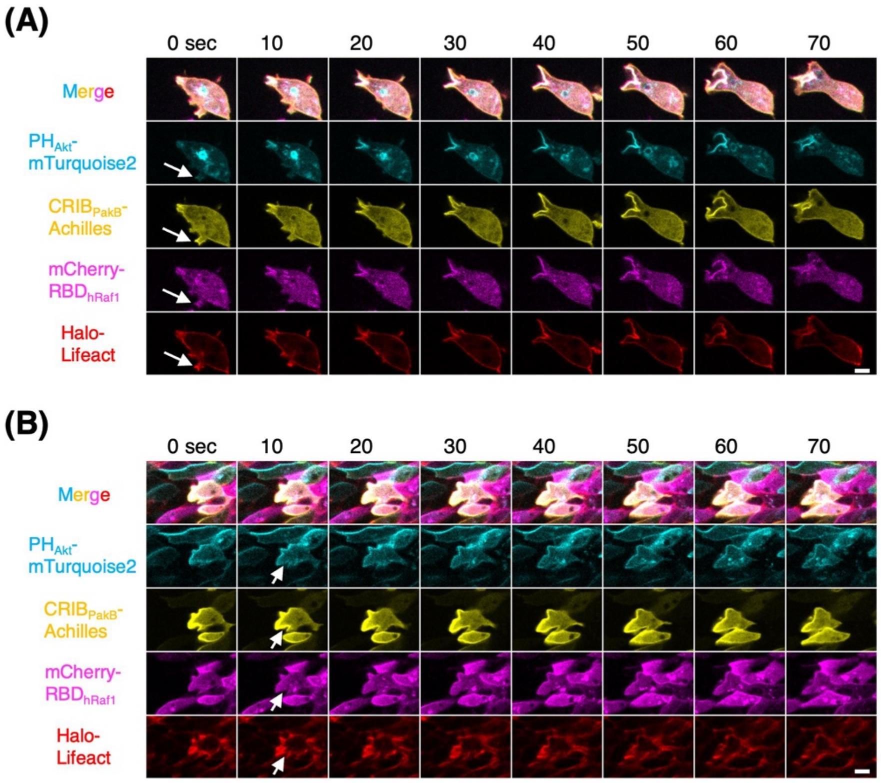


**Fig. S4. Application of four-color fluorescence imaging in *D. discoideum*.**

(A) Vegetative cells expressing actin related FP-tags. From top to bottom: merged channel, *coaAp*:PH$_{Akt}$-mTurquoise2 (PIP3; cyan), *coaAp*:CRIB$_{PakB}$-Achilles (Rac-GTP; green), *act15p*:mCherry-RBD$_{hRaf1}$ (Ras-GTP; magenta) and *act15p*:Halo-Lifeact (F-actin; yellow). HaloTag was labeled with Sarafluor650T. White arrows indicate protrusions with CRIB$_{PakB}$-Achilles and Halo-Lifeact. Scale bar, 10 µm. (B) Slug cells (posterior region) expressing the same set of FPs as in (A). The anterior-posterior axis of the slug is from left to right. White arrows: a protrusion with CRIB$_{PakB}$-Achilles and Halo-Lifeact. Scale bar, 10 µm.

| Fluorescent protein | Excitation (nm) | Emission (nm) | Application to live imaging of *D. discoideum* | Reference |
|---|---|---|---|---|
| **TagBFP** | 399 | 456 | (Kundert *et al.*, 2020) | (Subach *et al.*, 2008) |
| **mTagBFP2** | 399 | 454 | This study | (Subach *et al.*, 2011) |
| **LSSmGFP** | 400 | 510 | This study | (Campbell *et al.*, 2022) |
| **mTurquoise2** | 434 | 474 | (Mukai *et al.*, 2016) | (Goedhart *et al.*, 2012) |
| **GFP** | 395 | 509 | (Fey *et al.*, 1995) | (Chalfie *et al.*, 1994) |
| **GFP (S65T)** | 490 | 510 | (Aizawa *et al.*, 1997) | (Kilgard *et al.*, 1995) |
| **mNeonGreen** | 506 | 517 | (Antolović *et al.*, 2019) | (Shaner *et al.*, 2013) |
| **Achilles** | 513 | 525 | This study | (Yoshioka-Kobayashi *et al.*, 2020) |
| **mRFPmars** | 585 | 602 | (Fischer *et al.*, 2004) | (Fischer *et al.*, 2004) |
| **mCherry** | 587 | 610 | (Itoh and Yumura, 2007) | (Shaner *et al.*, 2004) |
| **tdTomato** | 554 | 581 | (Benabentos *et al.*, 2009) | (Shaner *et al.*, 2004) |
| **mScarlet** | 569 | 594 | (Paschke *et al.*, 2018) | (Bindels *et al.*, 2016) |
| **mScarlet-I** | 569 | 593 | This study | (Bindels *et al.*, 2016) |
| **iRFP** | 690 | 713 | (Ohta *et al.*, 2018) | (Filonov *et* |

| | | | | |
|---|---|---|---|---|
| | | | | *al.*, 2011) |
| **mIFP** | 683 | 704 | (Kundert *et al.*, 2020) | (Yu *et al.*, 2015) |
| **miRFP670nano3** | 645 | 670 | This study | (Oliinyk *et al.*, 2022) |
| **HaloTag** | – | – | (Matsuoka *et al.*, 2012) | (Los *et al.*, 2008) |

**Table S1. List of fluorescence protein used for imaging of *D. discoideum* cells.**

Gabriel, D., Hacker, U., Köhler, J., Müller-Taubenberger, A., Schwartz, J.M., Westphal, M. and

Gerisch, G. 1999. The contractile vacuole network of Dictyostelium as a distinct organelle: its dynamics visualized by a GFP marker protein. *J. Cell Sci.*, **112**: 3995–4005.
Goedhart, J., Von Stetten, D., Noirclerc-Savoye, M., Lelimousin, M., Joosen, L., Hink, M.A., Van Weeren, L., Gadella, T.W.J. and Royant, A. 2012. Structure-guided evolution of cyan fluorescent proteins towards a quantum yield of 93%. *Nat. Commun. 2012 31*, **3**: 1–9.
Hashimura, H., Morimoto, Y. V., Yasui, M. and Ueda, M. 2019. Collective cell migration of Dictyostelium without cAMP oscillations at multicellular stages. *Commun. Biol.*, **2**.
Itoh, G. and Yumura, S. 2007. A novel mitosis-specific dynamic actin structure in Dictyostelium cells. *J. Cell Sci.*, **120**: 4302–4309.
Kilgard, R., Heim, A.B. and Tsien, R.Y. 1995. Improved green fluorescence. *Nat. 1995 3736516*, **373**: 663–664.
Kundert, P., Sarrion-Perdigones, A., Gonzalez, Y., Katoh-Kurasawa, M., Hirose, S., Lehmann, P., Venken, K.J.T. and Shaulsky, G. 2020. A GoldenBraid cloning system for synthetic biology in social amoebae. *Nucleic Acids Res.*, **48**: 4139–4146.
Los, G. V., Encell, L.P., McDougall, M.G., Hartzell, D.D., Karassina, N., Zimprich, C., Wood, M.G., Learish, R., Ohana, R.F., Urh, M., Simpson, D., Mendez, J., Zimmerman, K., Otto, P., Vidugiris, G., Zhu, J., Darzins, A., Klaubert, D.H., Bulleit, R.F., *et al.* 2008. HaloTag: A novel protein labeling technology for cell imaging and protein analysis. *ACS Chem. Biol.*, **3**: 373–382.
Matsuoka, S., Miyanaga, Y., Yanagida, T. and Ueda, M. 2012. Single-Molecule Imaging of Stochastic Signaling Events in Living Cells. *Cold Spring Harb. Protoc.*, **2012**: pdb.top068189.
Meili, R., Ellsworth, C. and Firtel, R.A. 2000. A novel Akt/PKB-related kinase is essential for morphogenesis in Dictyostelium. *Curr. Biol.*, **10**: 708–717.
Meili, R., Ellsworth, C., Lee, S., Reddy, T.B., Ma, H., Firtel, R.A. and Hemmings, B.A. 1999. Chemoattractant-mediated transient activation and membrane localization of Akt/PKB is required for efficient chemotaxis to cAMP in Dictyostelium. *EMBO J.*, **18**: 2092–105.
Mukai, A., Ichiraku, A. and Horikawa, K. 2016. Reliable handling of highly A/T-rich genomic DNA for efficient generation of knockin strains of Dictyostelium discoideum. *BMC Biotechnol.*, **16**: 1–11.
Nakajima, A., Ishihara, S., Imoto, D. and Sawai, S. 2014. Rectified directional sensing in long-range cell migration. *Nat. Commun. 2014 51*, **5**: 1–14.
Ohta, Y., Furuta, T., Nagai, T. and Horikawa, K. 2018. Red fluorescent cAMP indicator with increased affinity and expanded dynamic range. *Sci. Rep.*, **8**: 1866.
Oliinyk, O.S., Baloban, M., Clark, C.L., Carey, E., Pletnev, S., Nimmerjahn, A. and Verkhusha, V. V. 2022. Single-domain near-infrared protein provides a scaffold for antigen-dependent fluorescent nanobodies. *Nat. Methods 2022 196*, **19**: 740–750.
Paschke, P., Knecht, D.A., Silale, A., Traynor, D., Williams, T.D., Thomason, P.A., Insall, R.H., Chubb, J.R., Kay, R.R. and Veltman, D.M. 2018. Rapid and efficient genetic engineering of both wild type and axenic strains of Dictyostelium discoideum. *PLoS One*, **13**: e0196809.
Perry, C.J., Warren, E.C., Damstra-Oddy, J.L., Storey, C., Francione, L.M., Annesley, S.J., Fisher, P.R., Müller-Taubenberger, A. and Williams, R.S.B. 2020. A Dictyostelium discoideum mitochondrial fluorescent tagging vector that does not affect respiratory function. *Biochem. Biophys. Reports*, **22**: 100751.
Riedl, J., Crevenna, A.H., Kessenbrock, K., Yu, J.H., Neukirchen, D., Bista, M., Bradke, F., Jenne, D., Holak, T.A., Werb, Z., Sixt, M. and Wedlich-Soldner, R. 2008. Lifeact: a versatile marker to visualize F-actin. *Nat. Methods 2008 57*, **5**: 605–607.
Rupper, A., Lee, K., Knecht, D. and Cardelli, J. 2001. Sequential activities of phosphoinositide 3-kinase, PKB/Akt, and Rab7 during macropinosome formation in Dictyostelium. *Mol. Biol. Cell*, **12**: 2813–2824.
Schneider, N., Schwartz, J.M., Köhler, J., Becker, M., Schwarz, H. and Gerisch, G. 2000. Golvesin-GFP fusions as distinct markers for Golgi and post-Golgi vesicles in Dictyostelium cells. *Biol. Cell*, **92**: 495–511.
Shaner, N.C., Campbell, R.E., Steinbach, P.A., Giepmans, B.N.G., Palmer, A.E. and Tsien, R.Y. 2004. Improved monomeric red, orange and yellow fluorescent proteins derived from

Discosoma sp. red fluorescent protein. *Nat. Biotechnol. 2004 2212*, **22**: 1567–1572.

Shaner, N.C., Lambert, G.G., Chammas, A., Ni, Y., Cranfill, P.J., Baird, M.A., Sell, B.R., Allen, J.R., Day, R.N., Israelsson, M., Davidson, M.W. and Wang, J. 2013. A bright monomeric green fluorescent protein derived from Branchiostoma lanceolatum. *Nat. Methods 2013 105*, **10**: 407–409.

Singleton, C.K., Manning, S.S. and Ken, R. 1989. Primary structure and regulation of vegetative specific genes of Dictyostelium discoideum. *Nucleic Acids Res.*, **17**: 9679.

Subach, O.M., Cranfill, P.J., Davidson, M.W. and Verkhusha, V. V. 2011. An Enhanced Monomeric Blue Fluorescent Protein with the High Chemical Stability of the Chromophore. *PLoS One*, **6**: e28674.

Subach, O.M., Gundorov, I.S., Yoshimura, M., Subach, F. V., Zhang, J., Grüenwald, D., Souslova, E.A., Chudakov, D.M. and Verkhusha, V. V. 2008. Conversion of Red Fluorescent Protein into a Bright Blue Probe. *Chem. Biol.*, **15**: 1116–1124.

Veltman, D.M., Akar, G., Bosgraaf, L. and Van Haastert, P.J.M. 2009. A new set of small, extrachromosomal expression vectors for Dictyostelium discoideum. *Plasmid*, **61**: 110–118.

Veltman, D.M., Keizer-Gunnink, I. and Haastert, P.J.M.V. 2009. An extrachromosomal, inducible expression system for Dictyostelium discoideum. *Plasmid*, **61**: 119–125.

Veltman, D.M., King, J.S., Machesky, L.M. and Insall, R.H. 2012. SCAR knockouts in Dictyostelium: WASP assumes SCAR's position and upstream regulators in pseudopods. *J. Cell Biol.*, **198**: 501–508.

Veltman, D.M., Williams, T.D., Bloomfield, G., Chen, B.C., Betzig, E., Insall, R.H. and Kay, R.R. 2016. A plasma membrane template for macropinocytic cups. *Elife*, **5**: 24.

Yoshioka-Kobayashi, K., Matsumiya, M., Niino, Y., Isomura, A., Kori, H., Miyawaki, A. and Kageyama, R. 2020. Coupling delay controls synchronized oscillation in the segmentation clock. *Nat. 2020 5807801*, **580**: 119–123.

Yu, D., Baird, M.A., Allen, J.R., Howe, E.S., Klassen, M.P., Reade, A., Makhijani, K., Song, Y., Liu, S., Murthy, Z., Zhang, S.Q., Weiner, O.D., Kornberg, T.B., Jan, Y.N., Davidson, M.W. and Shu, X. 2015. A naturally monomeric infrared fluorescent protein for protein labeling in vivo. *Nat. Methods 2015 128*, **12**: 763–765.

#### Supplemental movies

**Movie S1 Four-color imaging of mitochondria, plasma membrane, contractile vacuoles and nucleus in the vegetative cells (related to Fig. 10A)**

Fluorescent images of vegetative cells expressing 4-color markers. Fluorescence images of *act15p*:GcvH1(N99)-mTagBFP2 (cyan), *coaAp*:PKBR1(N150)-Achilles (yellow), *act15p*:Dajumin-mScarlet-I (magenta), *act15p*:HistoneH1-miRFP670nano3 (red) and merged images are shown. Images were acquired at 10 seconds intervals. Scale bar, 5 μm.

**Movie S2 Five-color imaging of mitochondria, cytoskeleton, plasma membrane, Golgi apparatus and nucleus in the slug cells (related to Fig. 10B)**

Fluorescent images of cells dissociated from slugs expressing the F-actin probe and organelle markers. Fluorescence images of *act15p*:GcvH1(N99)-mTagBFP2 (cyan, top middle), *act15p*:LSSmGFP-Lifeact (green, top right), *act15p*:Golvesin-Achilles (yellow, bottom left), *act5p*:PKBR1(N150)-mScarlet-I-2x (magenta, bottom middle), *act15p*:HistoneH1-miRFP670nano3 (red, bottom right), and the merged image (top left) were shown. Images were acquired at 10 seconds intervals. Scale bar, 10 μm.

**Movie S3 Four-color imaging of PIP3, Rac-GTP, Ras-GTP and F-actin in the vegetative cells (related to Fig. S4A)**

Fluorescent images of vegetative cells expressing 4-color markers. Fluorescence images of *coaAp*:$PH_{Akt}$-mTurquoise2 (cyan), *coaAp*:$CRIB_{PakB}$-Achilles (yellow), *act15p*:mCherry-$RBD_{hRaf1}$ (magenta), *act15p*:Halo-Lifeact (red), and merged images are shown. Images were acquired at 10 seconds intervals. Scale bar, 10 μm.

**Movie S4 Four-color imaging of PIP3, Rac-GTP, Ras-GTP and F-actin in the slug cells (related to Fig. S4B)**

Fluorescent images of vegetative cells expressing 4-color markers. Fluorescence images of *coaAp*:$PH_{Akt}$-mTurquoise2 (cyan), *coaAp*:$CRIB_{PakB}$-Achilles (yellow), *act15p*:mCherry-$RBD_{hRaf1}$ (magenta), *act15p*:Halo-Lifeact (red), and merged images are shown. Images were acquired at 10 seconds intervals. Scale bar, 10 µm.

**Movie S5 Cellulose imaging of extracellular matrix in the slug trail (related to Fig. 11)**

Fluorescence images of cellulose in the extracellular matrix labeled with calcofluor white. The left panel of the movie shows a bright-field image. The right panel shows fluorescence images of calcofluor white in agar used for labeling the cellulose in the slime sheath. Images were acquired at 20 seconds intervals. Scale bar, 10µm.
